# Software choice and depth of sequence coverage can impact plastid genome assembly – A case study in the narrow endemic *Calligonum bakuense*

**DOI:** 10.1101/2021.10.06.463392

**Authors:** Eka Giorgashvili, Katja Reichel, Calvinna Caswara, Vuqar Kerimov, Thomas Borsch, Michael Gruenstaeudl

**Author notes:** Corresponding author: Michael Gruenstaeudl.

## Abstract

Most plastid genome sequences are assembled from short-read whole-genome sequencing data, yet the impact that sequence coverage and the choice of assembly software can have on the accuracy of the resulting assemblies is poorly understood. In this study, we test the impact of both factors on plastid genome assembly in the threatened and rare endemic shrub *Calligonum bakuense,* which forms a distinct lineage in the genus *Calligonum.* We aim to characterize the differences across plastid genome assemblies generated by different assembly software tools and levels of sequence coverage and to determine if these differences are large enough to affect the phylogenetic position inferred for *C. bakuense.* Four assembly software tools (FastPlast, GetOrganelle, IOGA, and NOVOPlasty) and three levels of sequence coverage (original depth, 2,000x, and 500x) are compared in our analyses. The resulting assemblies are evaluated with regard to reproducibility, contig number, gene complement, inverted repeat length, and computation time; the impact of sequence differences on phylogenetic tree inference is also assessed. Our results show that software choice can have a considerable impact on the accuracy and reproducibility of plastid genome assembly and that GetOrganelle produced the most consistent assemblies for *C. bakuense.* Moreover, we found that a cap in sequence coverage can reduce both the sequence variability across assembly contigs and computation time. While no evidence was found that the sequence variability across assemblies was large enough to affect the phylogenetic position inferred for *C. bakuense,* differences among the assemblies may influence genotype recognition at the population level.

## INTRODUCTION

The comparative analysis of complete plastid genomes is performed in numerous investigations every year, even though the bioinformatic assembly of these genomes has not yet been perfected. Complete plastid genomes constitute a popular information source in various areas of plant evolutionary research, including phylogenetics (e.g., Xu et al., 2019; Koehler et al., 2020), phylogeography (e.g., Moner et al., 2018; del Valle et al., 2019), and population genetics (e.g., Yang et al., 2013; Rogalski et al., 2015). In recent years, the sequencing and comparison of dozens, if not hundreds, of complete plastid genomes per investigation has become commonplace (e.g., Saarela et al., 2018; Huang et al., 2019). Most of these studies generate complete plastid genomes from short-read whole-genome sequencing data (i.e., “genome skimming” data Bakker, 2017; Twyford and Ness, 2017). Several specialized software tools for the *de novo* assembly of plastid genomes from genome skimming data exist (e.g., Izan et al., 2017; McKain and Wilson, 2017; Coissac, 2017), but the process of generating complete and accurate assemblies from such data remains challenging (Wu et al., 2015; Freudenthal et al., 2020). For example, the use of genome skimming data for plastid genome assembly requires the separation of reads from different genomic compartments (Twyford and Ness, 2017). If done bioinformatically, this separation is only as accurate as the employed reference genome and its similarity to the target genome (Izan et al., 2017; Jin et al., 2020). Similarly, the use of genome skimming data often necessitates the assembly of reads that cover the target genome with unequal depth (Doorduin et al., 2011; Izan et al., 2017). An unequal sequence coverage runs contrary to the implicit assumption of many assembly algorithms that the input reads should cover the target genome homogeneously (Peng et al., 2012; McCorrison et al., 2014; Olson et al., 2019). Moreover, the quadripartite structure of plastid genomes, which consist of a long (LSC) and a short (SSC) single-copy region separated by two inverted repeats (IR) (Ruhlman and Jansen, 2014), often requires the manual circularization of linear assembly contigs (Twyford and Ness, 2017) because genome skimming data comprise an amalgamation of different reads, some of which support alternative junction sites (Jin et al., 2020). Furthermore, the direction of the SSC often needs to be homogenized across plastid genomes before their comparison due to the structural heteroplasmy of these genomes (Walker et al., 2015), and genome skimming data typically contain reads representing both configurations (Wang and Lanfear, 2019). Several software tools have been developed to accommodate some of these challenges (e.g., Ankenbrand et al., 2018; Carrion et al., 2020; Wu et al., 2021), but the process of plastid genome assembly from genome skimming data remains imperfect.

The choice of assembly software and the depth of sequence coverage have been highlighted as potential sources for low assembly quality among plastid genomes, but a characterization of their impact has yet to be conducted. Several recent investigations have reported factors that may influence the accuracy of plastid genome assembly, including software choice (Freudenthal et al., 2020) and sequence coverage (reviewed in Gruenstaeudl and Jenke, 2020). The choice of assembly software has been reported as a source of inconsistent genome assembly by several previous studies (e.g., Magoc et al., 2013; Morrison et al., 2014). In the de novo assembly of plastid genomes from genome skimming data, these findings may be associated with different assembly algorithms: while some software tools have implemented algorithms that conduct a cyclical sequence extension from a single “seed” sequence (e.g., Dierckxsens et al., 2017), others employ a kmer-based construction of contigs, followed by the concatenation of multiple contigs based on sequence overlap and similarity to a reference genome (e.g., Bakker et al., 2016; McKain and Wilson, 2017). Accordingly, Freudenthal et al. (2020) found considerable differences among the results of different assembly software despite employing the same input sequence data. Interestingly, many of the assembly differences identified by Freudenthal et al. (2020) corresponded to competing locations or orientations of the four plastid genome regions rather than nucleotide polymorphisms. The question if alternative plastid genome assemblies generated for the same taxon would impact a given downstream analysis such as species identification or phylogenetic inference has so far not been addressed. Differences in sequence coverage have also been reported as a source for distinct plastid genome assemblies. Doorduin et al. (2011), for example, found that the number of SNPs across the plastid genomes of multiple individuals of *Jacobaea vulgaris* varied between different regions of the genome depending on the depth of sequence coverage. Similarly, Kim et al. (2015) reported a correlation between cases of local mis-assembly and regions with exceptionally high sequence coverage in plastid genomes of rice; regions of exceptionally high coverage depth often occur when employing genome skimming data (Twyford and Ness, 2017). Moreover, Izan et al. (2017) found that regions with low sequence coverage were not correctly assembled under default software settings in several angiosperm plastid genomes. Indeed, genome assemblies with an unequal depth of sequence coverage are often characterized by high rates of sequencing error (Hubisz et al., 2011). These observations have rendered sequence coverage a popular indicator for assembly quality, especially in plastid genomes (Gruenstaeudl and Jenke, 2020). Gu et al. (2016), for example, employed sequence coverage as an indicator to refine the assembly of the plastid genome of *Lagerstroemia fauriei.* Despite the importance of sequence coverage for the successful assembly of plastid genomes, few, if any, studies have aimed to characterize the resulting assembly differences or evaluated if those differences are large enough to impact downstream analyses.

In this study, we test the impact of software choice and sequence coverage on the process of plastid genome assembly in a species for which a correct assembly is vital for conservation efforts. Specifically, we use the threatened and narrow endemic shrub *Calligonum bakuense* (Polygonaceae) as a test case for evaluating the variability in plastid genome assembly caused by software choice and coverage depth. The entire species comprises only 170–200 individuals which are currently inhabiting approximately seven localities around the Absheron Peninsula near Baku, the capital city of the Republic of Azerbaijan. *Calligonum bakuense* represents an exemplary case where a precise assembly of the plastid genome is of great importance to delineate the species, determine its correct phylogenetic placement in relation to other members of the genus, and assess its genetic diversity at the population level. Genomic information on *C. bakuense* is currently entirely absent, and documenting its complete plastid genome would be an important asset for future investigations on this rare and declining species. In this study, we use genome skimming data from two individuals of *C. bakuense* to characterize differences across genome assemblies in response to the choice of assembly software and levels of sequence coverage. Specifically, we test whether the plastid genome assembly of *C. bakuense* is consistent across four commonly employed assembly software tools and three different levels of sequence coverage, and if any differences among the resulting assemblies can potentially affect the outcome of phylogenetic tree inference. Based on our findings, we discuss the consequences that differences in plastid genome assembly of the magnitude detected here could have on biological conclusions and we make recommendations to optimize the assembly of complete plastid genomes.

## MATERIALS AND METHODS

### Biology and distribution of *Calligonum bakuense*

*Calligonum bakuense* LITV. is a psammophytic shrub endemic to coastal sand dune areas along the western Caspian shoreline near the city of Baku (Karjagin, 1952; Soskov and Akhmed-Zade, 1974). The species is a unique and declining element of the flora of Azerbaijan and of high conservation interest (Atamov, 2008). It currently comprises a total of seven wild populations that are distributed across a distance of approximately 120 km and collectively contain roughly 170–200 individuals (Figure 1). Here we assemble and report the plastid genomes of two individuals that represent the northern- and the southernmost localities of its current distribution area. The evolutionary relationships of *C. bakuense* to other members of *Calligonum* are currently unknown, as is the population structure within the species. *Calligonum* L. is a lineage of xerophytic shrubs with an estimated 30–40 species; it is distributed from northern Africa, the Arab Peninsula, South West Asia, the Caucasus, the Irano-Turanian region, and Central Asia to China (Brandbyge, 1993; Abdellaoui et al., 2011). Several species of the genus are globally red-listed and exhibit declining population sizes (Baillie et al., 2004). Currently, there is no comprehensive molecular phylogeny of *Calligonum,* but complete plastid genome sequences have been shown as a promising basis for inferring phylogenetic relationships among Chinese members of the genus (Song et al., 2020).

**Figure 1.**
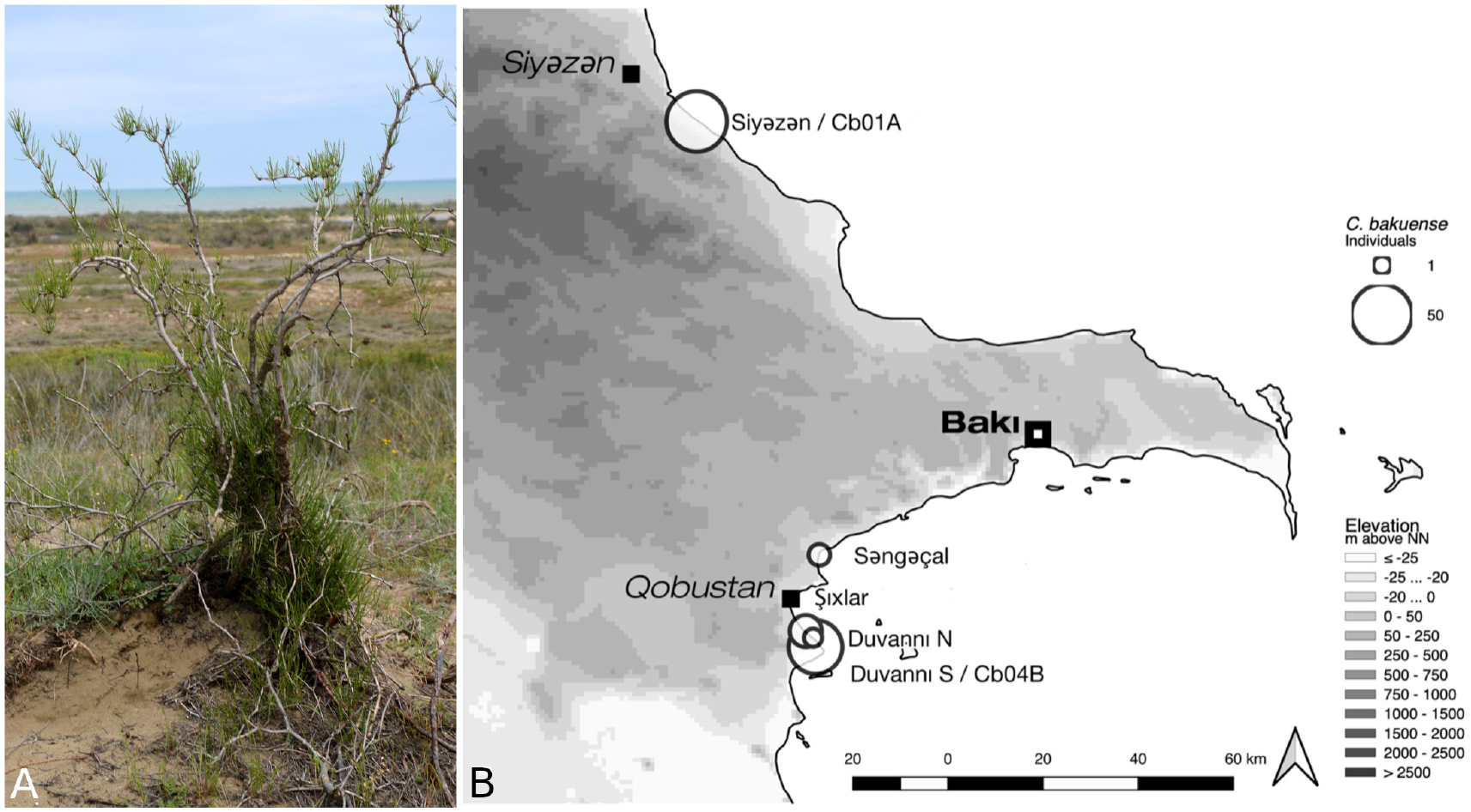
Habit and natural environment (A) and the current distribution area (B) of *C. bakuense.* The distribution map indicates the localities of all sampled natural populations of *C. bakuense,* including the localities that Cb01A and Cb04B were sampled from.

### DNA extraction and genome skimming

In preparation for a population genetic study on *C. bakuense,* silica-dried tissue samples from all known individuals of the species were collected during several field trips between 2013 and 2015. A plant individual (Cb01A) that represents the northernmost and an individual (Cb04B) that represents the southernmost locality of the current distribution area were selected for low-coverage whole-genome sequencing (Figure 1). Whole genomic DNA of each individual was extracted using a modified CTAB protocol (Borsch et al., 2003), sheared via ultrasonication to an average fragment size of ~300 bp, and converted to a barcoded genomic library using the Illumina TruSeq DNA sample preparation kit (Illumina, San Diego, CA, USA) under the high sample protocol of the manufacturer. The DNA of both individuals was pooled equimolarly and then sequenced on a full Illumina HiSeq 4000 plate by Macrogen Inc. (Seoul, South Korea). After sequencing, low-quality bases (phred-score <20) and any remnants of Illumina adapter sequences were trimmed from the raw reads with Cutadapt v. 1.14 (Martin, 2011).

### Bioinformatic extraction of plastid genome reads

Conducting the plastid genome assembly of *C. bakuense* on the raw set of sequence reads generated via genome skimming would have exceeded the maximum capacity of some of the assembly software tools under study. Consequently, we extracted those reads of the genome skimming dataset prior to genome assembly that represent the plastid genome. Specifically, we mapped the raw sequence reads to a set of related, previously published plastid genomes and then bioinformatically extracted and retained only the successfully mapped reads. To avoid any bias in this read extraction, we selected complete plastid genomes of twelve different genera of the plant order Caryophyllales as reference genomes; half of these genomes represent the family Polygonaceae. These reference genomes were: *Fagopyrum esculentum* subsp. *ancestrale* (GenBank accession number NC 010776), *Fallopia multiflora* (NC_041239), *Rumex acetosa* (NC_042390), *Muehlenbeckia australis* (MG604297), *Oxyria sinensis* (NC_032031), and *Rheum palmatum* (NC_027728, all Polygonaceae); *Amaranthus hypochondriacus* (NC_030770) and *Chenopodium quinoa* (NC_034949, both Amaranthaceae); *Mesembryanthemum crystallinum* (NC_029049, Aizoaceae); *Carnegiea gigantea* (NC 027618, Cactaceae); *Dianthus caryophyllus* (NC 039650, Caryophyllaceae); and *Nyctaginia capitata* (NC_041415, Nyctaginaceae). The process of read mapping and extraction was conducted using script 5 of the pipeline described in Gruenstaeudl et al. (2018).

### Capping of sequence coverage

To determine if the process of plastid genome assembly is consistent across different levels of sequence coverage, we created subsets (hereafter “capped read sets”) of the complete plastid genome read set with a lower average sequence coverage. Specifically, the depth of sequence coverage was capped at 2,000x and at 500x, which represents approximately 25% and 6.25%, respectively, of the average uncapped sequence coverage. This cap was not a hard threshold above which all additional reads were removed, but a soft threshold above which additional reads were progressively curtailed: we employed a one-tailed normalization of sequence coverage depth using the script ‘bbnorm.sh’ of the software BBtools v.33.89 (Bushnell, 2015) under default settings and using the plastid genome of *Calligonum caput-medusae* (MN202600; Song et al., 2020) as a structural reference. The capped read sets were treated identically to the complete read set during genome assembly and in all subsequent analyses.

### Genome assembly

To determine if the process of plastid genome assembly for *C. bakuense* is consistent across different assembly software tools, we employed and compared the assembly results of four different tools: NOVO-Plasty v.3.8.3 (Dierckxsens et al., 2017), GetOrganelle v.1.6.4 (Jin et al., 2020), FastPlast v.1.2.8 (McKain and Wilson, 2017), and IOGA v.38.26 (Bakker et al., 2016). Each of these software tools had been designed for the *de novo* assembly of plastid genomes from short sequence reads and had demonstrated its utility in previous plastid genomic studies (reviewed in Freudenthal et al., 2020). To improve the comparability of the assembly process across these tools, we employed each software under its default settings. To ensure a uniform software execution and to compare computation times across the tools, all assemblies of *C. bakuense* were conducted on the high-performance computer cluster ‘Curta’ of the Freie Universitat Berlin under the following settings: a single 64-bit processor, an allotment of 2 GB of RAM, and a disk I/O speed of 129 MB/s. The raw output as well as the log file of each assembly software run are available on Zenodo under https://zenodo.org/record/3999863.

In practice, plastid genome assembly software often generates multiple incomplete, linear contigs instead of a complete, circular genome sequence (Twyford and Ness, 2017). Incomplete contigs typically require manual intervention to be combined into a complete genome sequence (Gruenstaeudl et al., 2018). In this study, two of the software tools produced incomplete contigs for *C. bakuense*. We concatenated the incomplete contigs upon removing any end overhangs, followed by a circularization of the resulting super-contig. The concatenation of contigs was conducted by hand in Geneious v.11.1.4 (Kearse et al., 2012) through aligning each contig to the structural reference genome (*C. caput-medusae*) and then sorting the contigs according to their relative position. If adjacent contigs overlapped for at least 15 bp without differences in their nucleotide sequence, they were merged into a larger contig until all such contigs were combined into a single super-contig.

The identification of the endpoint of a circular genome sequence is challenging for most genome assembly algorithms (but see Wu et al., 2021) and often results in the detection of different endpoints across tools. To avoid inflating the number of differences between assemblies due to unequal endpoints, we manually corrected super-contigs if the inferred endpoints were within 100 bp across assemblies. Specifically, we searched for the first and the last 25 bp of the super-contig of each assembly via separate motif searches, with the maximum number of mismatches set to three bp. Any matches within 100 bp of the super-contig ends were considered to be instances where the assembly process extended the sequence beyond the actual endpoint. Such sequence motifs were removed from one of the two ends, followed by circularization of the super-contig. Similarly, poly-N motifs in contigs are often generated by plastid genome assembly software to indicate areas of sequence uncertainty. To avoid inflating the number of differences between assemblies, we automatically corrected poly-N-motifs using the software Pilon v.1.23 (Walker et al., 2014). Moreover, most software tools for plastid genome assembly do not automatically standardize the orientation of the SSC across assemblies, even though plastid genome isomers with alternative SSC orientations naturally exist in most land plants (Walker et al., 2015). To avoid inflating the number of differences between assemblies, we manually homogenized the orientation of the SSC across assemblies using Geneious.

### Replication of assembly runs

Several software tools for plastid genome assembly constitute bioinformatic pipelines rather than single applications (Gruenstaeudl et al., 2018). These pipelines typically employ a third-party assembly tool as their core assembly engine to conduct a k-mer based alignment of reads for the inference of de Bruijn graphs (Izan et al., 2017). FastPlast, IOGA, and GetOrganelle, for example, utilize the assembly software SPAdes (Bankevich et al., 2012) as core assembler, even though the full reproducibility of bacterial genome assemblies with SPAdes has been called into question (e.g., Liao et al., 2015; Souvorov et al., 2018). To characterize potential occurrences of spurious, non-reproducible inferences of de Bruijn graphs, we conducted every plastid genome assembly of the uncapped read set twice under the same input data and software settings (i.e., replicate run #1 and #2). The comparison of these replicate runs allowed us to ascertain the baseline replicability of plastid genome assemblies under these assembly tools.

The cyclical sequence extension that starts from a single “seed” sequence and is implemented in several assembly algorithms (e.g., Dierckxsens et al., 2017) may represent a source of contig variability not present in other assembly algorithms. Several studies have reported minor differences in the number and sequence of assembly contigs depending on the precise seed sequence employed and have, therefore, attempted to identify universally applicable seed sequences (e.g., Lim et al., 2018; Wu et al., 2021). To ensure that seed selection did not inflate the number of differences between assemblies, we employed the same seed sequence for each plastid genome assembly with NOVOPlasty. Moreover, we evaluated if seed selection represented a relevant source of contig variability in our dataset and, thus, duplicated all plastid genome assemblies under NOVOPlasty using a second seed sequence. Both seeds (i.e., seed #1 and seed #2) were arbitrarily selected from the read set.

### Sequence annotation

To enable consistent sequence annotations across all plastid genome assemblies of *C. bakuense,* the sequence annotations from two existing plastid genomes of *Calligonum* were transferred to the new assemblies using Geneious. Specifically, we automatically transferred all gene, tRNA, and rRNA annotations from the plastid genomes of *C. caput-medusae* and *C. arborescens* (MN202599; both Song et al., 2020) to the assemblies of *C. bakuense* based on a sequence similarity threshold of 95%. Upon transfer, we conducted a manual inspection of the transferred annotations for each coding region regarding the presence of start and stop codons, the absence of internal stop codons, and their lengths as a multiple of three. Any premature stop codon that was introduced by the transfer process but not based on the nucleotide sequence was corrected; any premature stop codon based on the nucleotide sequence was recorded as an indicator of low assembly quality (Table 2). The annotations of the IRs and, by extension, of the single copy regions were inferred for each assembly using script 4 of the pipeline of Gruenstaeudl et al. (2018).

### Evaluation of assembly quality

To assess the quality of the plastid genome assemblies of *C. bakuense* and, simultaneously, the performance of each assembly software, the raw output of each assembly process was evaluated with Quast v.4.6.3 (Gurevich et al., 2013). Specifically, we assessed and compared the number, length, and contiguity of the contigs generated by each assembly software. NGA50 and LGA50 were calculated as contiguity metrics (Earl et al., 2011; Gurevich et al., 2013). As part of this quality assessment, we also compared the computation times of the different assembly software tools. All assembly statistics were calculated after the removal of contigs smaller than 100 bp, if any, to avoid the counting of mono- or di-nucleotide fragments.

### 0.1 Characterization of assembly differences

To compare the different plastid genome assemblies of *C. bakuense* as generated by different software tools, levels of sequence coverage, seed selection, and run replication, we conducted a series of statistical evaluations based on pairwise genetic distances. As the basis for these comparisons, we generated all-against-all pairwise alignments of the assemblies using MAFFT v.7.471 (Katoh and Standley, 2013) under default settings. We then inferred the differences in sequence as well as in length of the four genome regions (i.e., LSCs, IRb, SSCs, and IRa) for each plastid genome pair. Sequence differences were calculated as the number of single nucleotide polymorphisms (SNPs) when excluding gaps but including nucleotide ambiguities using trimAl v.1.2 (Capella-Gutierrez et al., 2009). Upon calculation, difference values were aggregated in a pairwise genetic distance matrix. Since only the plastid genomes assembled with GetOrganelle were identical across the tested parameters, we designated the assemblies inferred with GetOrganelle on the 500x capped read set as the “final” plastid genome sequence for each individual under study. To visualize the genetic distances among all assemblies and both plant individuals, we conducted principal coordinates analyses (PCoAs). When visualizing the PCoA results, we centered the result projections on the values for the final plastid genome sequences (i.e., the assemblies generated with GetOrganelle for the read set capped at 500x), plotted the first two coordinates of results under a linear scale ranging from −1 to 1, and displayed the absolute variance (in bp) and the percentage of total variance along each axis within the plot. Since PCoAs can potentially distort pairwise distances between data points, we also plotted an overview of the genetic distances as caused by changes in software (including seed selection) and coverage cap (including run replication). Moreover, we also conducted PCoAs between the two plant individuals to have a biologically meaningful standard for the assembly differences within each individual. To that end, we calculated and plotted the pairwise genetic distances between Cb01A (set as origin) and Cb04B across software, coverage depth, seed selection, and run replicate. All calculations and visualizations based on the pairwise genetic distance matrices were conducted in R v.4.0.0 (R Development Core Team, 2019).

### Visualizations of region length, sequence coverage, and SNP location

Three types of visualization were employed to illustrate the structural and sequence differences between the plastid genome assemblies of *C. bakuense* as generated under different assembly parameters. First, we illustrated the differences in length between the LSC, the SSC, and the two IRs across the assemblies through an alignment overview of the four plastid genome regions using Geneious. Second, we visualized the depth of sequence coverage across the entire plastid genome sequence and in relation to the position of the genes and the four genome regions with PACVr v.1.0 using a calculation window of 250bp (Gruenstaeudl and Jenke, 2020). Third, we determined and visualized the location of SNPs between the plastid genome assemblies and in relation to changes in sequence coverage through pairwise comparisons of each assembly to the final genome sequence using MAFFT for sequence alignment and trimAl for SNP detection. We also visualized SNP locations in relation to the position of the genes and the three regions of the plastid genome using ShinyCircos v.29052020 (Yu et al., 2018).

### Phylogenetic inference

To test if the sequence differences among the plastid genome assemblies of *C. bakuense* are large enough to affect the phylogenetic placement of *C. bakuense* within *Calligonum*, we inferred the phylogenetic position of all plastid genome assemblies generated in this study within a large set of plastid genomes of *Calligonum.* Specifically, we retrieved all available plastid genomes of *Calligonum* that were stored on NCBI GenBank as of 30-Nov-2020 and combined these 21 genome records with the 40 genome assemblies of *C. bakuense* generated here with different software tools, levels of sequence coverage depth, seed sequences, and run replicates. We extracted and aligned all 81 protein-coding regions from each of the 61 genome records using script 9 of Gruenstaeudl et al. (2018) and inferred the best tree under the maximum likelihood (ML) criterion using RAxML v.8.2.9 (Stamatakis, 2014). To infer the actual phylogenetic position of *C. bakuense* among other species of *Calligonum*, we also conducted a second, more specific phylogenetic reconstruction. Specifically, we combined the 21 genome records of *Calligonum* stored on NCBI GenBank with only the two final plastid genome sequences of *C. bakuense,* added the plastid genome of *Rheum palmatum* (GenBank accession KR816224) as an outgroup, extracted and aligned all 81 protein-coding regions from this sequence set, and inferred the best ML tree using RAxML. *Rheum palmatum* was also used as outgroup in the study of Song et al. (2020). For both tree reconstructions, clade support was inferred through 100 bootstrap (BS) replicates generated under the rapid BS algorithm.

## RESULTS

### Number of sequence reads

Genome skimming of the two individuals of *C. bakuense* resulted in a total of 151,567,745 paired raw sequence reads for Cb01A and a total of 166,362,653 paired raw sequence reads for Cb04B. Upon extraction of the plastid genome reads, we counted 5,062,912 paired reads (3.3% of raw reads) for Cb01A and 2,998,391 paired reads (1.8%) for Cb04B. Upon capping sequence coverage, the read sets with a coverage depth of 2,000x exhibited 1,181,510 paired reads (0.78% of raw reads) for Cb01A and 1,149,375 paired reads (0.69%) for Cb04B; the capped read sets with a coverage depth of 500x exhibited 332,662 paired reads (0.22%) for Cb01A and 333,839 paired reads (0.20%) for Cb04B.

### Impact of software choice

The choice of assembly software had a considerable effect on the number and size of the generated assembly contigs, the contiguity of the assemblies, sequence equality of the inferred IRs, and the time required to conduct each assembly (Table 1). While some software tools assembled the plastid genome of *C. bakuense* as a single contig, others did not. GetOrganelle und IOGA represented the extremes among the tested software tools: GetOrganelle succeeded in assembling the complete plastid genome as a single contig under nearly all settings, whereas IOGA failed in this task under all settings. Under the original sequence coverage depth, GetOrganelle assembled the complete plastid genome of *C. bakuense* into a single, circular contig for both individuals and run replicates, precluding the need for any manual post-processing of the assembly contigs. Similarly, NOVOPlasty succeeded in assembling the complete plastid genome of *C. bakuense* as a single, circular contig under the original coverage depth for both individuals, run replicates, and seed sequences. For Cb01A, however, the assemblies generated with NOVOPlasty exhibited considerable size variability and often exceeded the length of the final plastid genome sequence; moreover, the inferred IRs were not identical in one of the assemblies. FastPlast also succeeded in assembling the complete plastid genome of *C. bakuense* as a single, circular contig under the original coverage depth. However, the contigs produced for both individuals and both replicate runs lagged or exceeded the length of the final plastid genome sequences due to incomplete or duplicated sections of the IRs, ranging from 201 kb to 143 kb in Cb01A and from 192 kb to 175 kb in Cb04B. The smaller than expected contig lacked a section of the IRa, whereas the larger than expected contigs exhibited a duplication of sections of the LSC adjacent to the IRs, necessitating manual post-processing of the assembly contigs and affecting the calculation of NGA50. IOGA, by contrast, did not succeed in assembling the complete plastid genome of *C. bakuense* as a single, complete contig under any setting. For both individuals, it generated more than 20 separate contigs, which represented only sections of the complete genome. Hence, the IOGA contigs had to be manually concatenated for both individuals and run replicates to generate circular assemblies. Moreover, the contigs assembled by IOGA for Cb01A did not imply identical IRs in one run replicate, indicating further issues. Computation times differed strongly across software tools and – in the case of IOGA and FastPlast – across run replicates but were similar across different seed sequences in NOVOPlasty. Under the original sequence coverage depth, GetOrganelle and NOVOPlasty were typically the quickest to generate assembly contigs, whereas FastPlast and IOGA often required a multiple of their computation time.

**Table 1.** Assembly statistics for the plastid genomes of the two individuals of *C. bakuense* under study. The evaluated assemblies were generated under different assembly software tools, sequence coverage depths, seed sequences, and run replicates. The assemblies that represent the final genome sequences are highlighted in bold. The four software tools employed are abbreviated as ‘FaPl’ (for FastPlast), ‘GetO’ (for GetOrganelles), ‘IOGA’, and ‘NOVO’ (for NOVOPlasty). The read sets are abbreviated as ‘repl1’ or ‘repl2’ for replicate run 1 or 2, respectively. Read sets with the original sequence coverage depth are abbreviated as ‘orig.’. Abbreviations used: asmb. = assembly; comp. = computation; cov. = coverage; equal. = equality in sequence; repl. = replicate.

| Asmb. | Cov. | Repl. | NOVO seed | Contigs | Largest contig (bp) | NGA50 (bp) | LGA50 | IR equal. | Comp. time (h, min.) |
| --- | --- | --- | --- | --- | --- | --- | --- | --- | --- |
| *Cb01A* |  |  |  |  |  |  |  |  |  |
| FaPl | orig. | repl1 |  | 1 | 200,694 | 118,168 | 1 | No | 05 h 20 min |
| FaPl | orig. | repl2 |  | 1 | 143,261 | 135,202 | 1 | No | 06 h 40 min |
| FaPl | 2000x |  |  | 1 | 162,404 | 162,128 | 1 | Yes | 01 h 16 min |
| FaPl | 500x |  |  | 1 | 163,292 | 162,896 | 1 | Yes | 24 min |
| GetO | orig. | repl1 |  | 1 | 162,128 | 162,128 | 1 | Yes | 44 min |
| GetO | orig. | repl2 |  | 1 | 162,128 | 162,128 | 1 | Yes | 44 min |
| GetO | 2000x |  |  | 2 | 118,241 | 118,215 | 1 | Yes | 01 h 08 min |
| **GetO** | **500x** |  |  | **1** | **162,128** | **162,128** | **1** | **Yes** | **20 min** |
| IOGA | orig. | repl1 |  | 21 | 89,039 | 88,068 | 1 | Yes | 09 h 42 min |
| IOGA | orig. | repl2 |  | 21 | 89,039 | 88,068 | 1 | No | 06 h 43 min |
| IOGA | 2000x |  |  | 83 | 129,550 | 118,520 | 1 | No | 07 h 50 min |
| IOGA | 500x |  |  | 51 | 91,976 | 89,718 | 1 | No | 02 h 22 min |
| NOVO | orig. | repl1 | seed1 | 1 | 170,093 | 131,660 | 1 | No | 01 h 05 min |
| NOVO | orig. | repl2 | seed1 | 1 | 170,099 | 170,099 | 1 | Yes | 57 min |
| NOVO | 2000x |  | seed1 | 1 | 162,128 | 162,128 | 1 | Yes | 23 min |
| NOVO | 500x |  | seed1 | 1 | 162,128 | 162,128 | 1 | Yes | 07 min |
| NOVO | orig. | repl1 | seed2 | 1 | 162,128 | 162,128 | 1 | Yes | 01 h 05 min |
| NOVO | orig. | repl2 | seed2 | 1 | 170,106 | 170,106 | 1 | Yes | 01 h 00 min |
| NOVO | 2000x |  | seed2 | 1 | 162,128 | 162,128 | 1 | Yes | 23 min |
| NOVO | 500x |  | seed2 | 1 | 162,128 | 162,128 | 1 | Yes | 07 min |
| *Cb04B* |  |  |  |  |  |  |  |  |  |
| FaPl | orig. | repl1 |  | 1 | 175,272 | 175,272 | 1 | Yes | 09 h 24 min |
| FaPl | orig. | repl2 |  | 1 | 192,943 | 118,215 | 1 | No | 03 h 36 min |
| FaPl | 2000x |  |  | 1 | 163,890 | 162,129 | 1 | Yes | 01 h 16 min |
| FaPl | 500x |  |  | 1 | 163,292 | 163,292 | 1 | Yes | 24 min |
| GetO | orig. | repl1 |  | 1 | 162,129 | 162,129 | 1 | Yes | 04 h 16 min |
| GetO | orig. | repl2 |  | 1 | 162,129 | 162,129 | 1 | Yes | 01 h 04 min |
| GetO | 2000x |  |  | 2 | 118,238 | 118,215 | 1 | Yes | 01 h 28 min |
| **GetO** | **500x** |  |  | **1** | **162,129** | **162,129** | **1** | **Yes** | **20 min** |
| IOGA | orig. | repl1 |  | 54 | 90,241 | 88,240 | 1 | Yes | 11 h 14 min |
| IOGA | orig. | repl2 |  | 85 | 90,630 | 87,507 | 1 | Yes | 07 h 42 min |
| IOGA | 2000x |  |  | 102 | 55,966 | 27,790 | 2 | No | 08 h 17 min |
| IOGA | 500x |  |  | 40 | 75,285 | 74,394 | 1 | No | 03 h 18 min |
| NOVO | orig. | repl1 | seed1 | 1 | 162,129 | 162,129 | 1 | Yes | 01 h 23 min |
| NOVO | orig. | repl2 | seed1 | 1 | 162,129 | 162,129 | 1 | Yes | 01 h 24 min |
| NOVO | 2000x |  | seed1 | 1 | 162,129 | 162,129 | 1 | Yes | 20 min |
| NOVO | 500x |  | seed1 | 1 | 162,129 | 162,129 | 1 | Yes | 06 min |
| NOVO | orig. | repl1 | seed2 | 1 | 162,129 | 162,129 | 1 | Yes | 01 h 00 min |
| NOVO | orig. | repl2 | seed2 | 1 | 162,129 | 162,129 | 1 | Yes | 01 h 32 min |
| NOVO | 2000x |  | seed2 | 1 | 162,129 | 162,129 | 1 | Yes | 20 min |
| NOVO | 500x |  | seed2 | 1 | 162,129 | 162,129 | 1 | Yes | 06 min |

### Impact of sequence coverage depth

The depth of sequence coverage also had a considerable effect on the number and size of the generated assembly contigs, the contiguity of the assemblies, sequence equality of the inferred IRs, and the time required to conduct each assembly (Table 1). For example, we observed that GetOrganelle assembled the complete plastid genome of *C. bakuense* into a single, circular contig under the original coverage depth and a depth of 500x. For a coverage depth of 2,000x, however, GetOrganelle generated two contigs, which had to be concatenated to create a complete genome sequence. The break point between the two contigs was located at the junction site between IRb and the SSC in both individuals, indicating that this non-contiguity was correlated with the quadripartite genome structure. NOVOPlasty also appeared to be largely insensitive to changes in coverage depth and generated identical genome sequences for Cb04B and a single, circular contig with occasional length differences for Cb01A across different depth levels. Accordingly, the largest contig retrieved for each individual of *C. bakuense* under a coverage depth of 500x was identical under NOVOPlasty and GetOrganelle and represented the final plastid genome sequence of each individual. Under FastPlast, by contrast, all genome assemblies were different in length for different coverage depths. Moreover, the IRs of the assembled plastid genomes were found to be identical within assemblies only under the capped read sets as well as replicate run 1 of the uncapped read set in Cb04B. The assembly process conducted by IOGA appeared to be even more sensitive to changes in coverage depth: for individual Cb01A, IOGA assembled 21 contigs under the original read set, 83 contigs under a coverage cap of 2,000x, and 51 contigs under a coverage cap of 500x; for Cb04B, the software generated between 54 and 85 contigs under the original read set (depending on the run replicate), 102 contigs under a coverage cap of 2,000x, and 40 contigs under a coverage cap of 500x. While at least half of the final genome sequence was encompassed within a single contig in all but one of these cases, the assembly results for each coverage depth level had to be manually concatenated to generate complete plastid genomes. Computation times differed strongly across different depths of sequence coverage and generally correlated with the size of the input dataset: datasets with a capped sequence depth were typically analyzed faster than the uncapped datasets. NOVOPlasty was the software that achieved a complete plastid genome assembly for *C. bakuense* in the shortest amount of time if the sequence coverage was capped at 500x.

In summary, we found that among the four assembly software tools tested, only GetOrganelle generated genome assemblies that were identical in both length and sequence across all replicates and levels of sequence coverage depth within an individual. The split of the assembly sequence by GetOrganelle into two separate contigs under a coverage cap of 2,000x did not invalidate this observation, as the break point was located exactly at the junction between IRb and the SSC, which is a natural break point in a circular quadripartite genome. We, therefore, considered the sequences generated with GetOrganelle for the two individuals of *C. bakuense* as the best results and submitted them as the official plastid genome sequences of the species to GenBank (accessions MT806099 and MT806098 for Cb01A and Cb04B, respectively; Figure 2). Under this assumption, the plastid genomes of Cb01A and Cb04B are almost identical and differ only by a single nucleotide: one adenine within a poly-A microsatellite located in the intergenic spacer between the genes *ndhF* and *rpl32* of the SSC region of the genome. Plastid genome diversity within *C. bakuense* is, thus, extremely low, but not zero.

**Figure 2.**
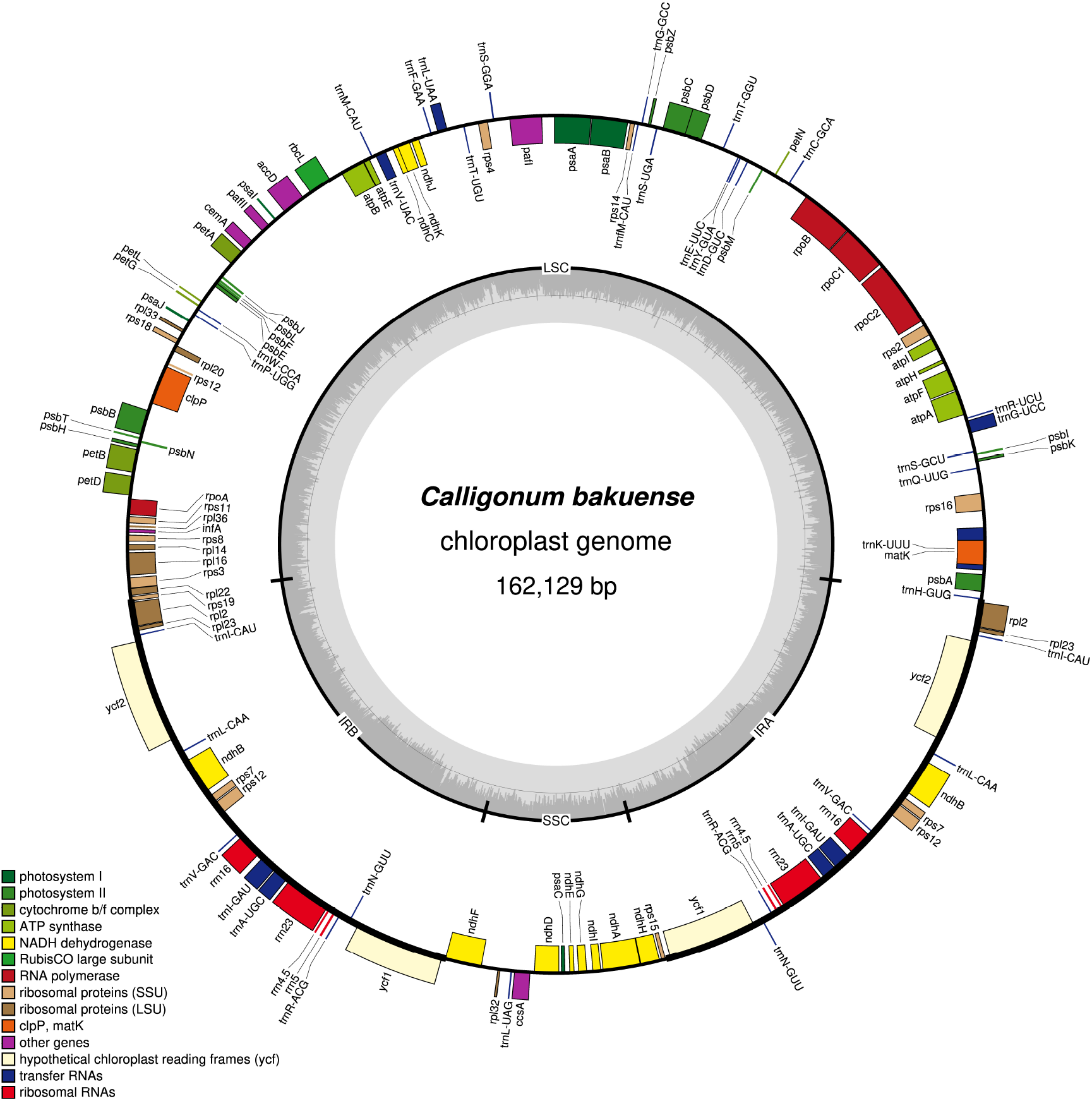
Map of the complete plastid genome of the plant individual Cb01A of *C. bakuense* as assembled by GetOrganelle under a coverage cap of 500x. This assembly represents the final plastid genome sequence for Cb01A.

### Characterization of assembly differences

The characterization of the number of SNPs and the length differences within each of the four plastid genome regions in relation to different software tools and depths of sequence coverage through PCoA indicated the presence of a complex pattern of differences (Figure 3A). For example, the different assembly software tools produced assemblies that were heterogeneous in both length and sequence for at least one of the two individuals. The PCoA plots indicated no single parameter that explained the deviation across all software tools or levels of coverage depth. For the lengths of the four plastid genome regions of Cb04B, the first coordinate of the PCoA explained nearly all variance in the dataset, indicating the presence of only one or two nearly identical outlier assemblies; this pattern was not observed for the LSC and IR length under Cb01A, for which the greater number of outliers is reflected in a reduced explained variance for the first two coordinates. For the number of SNPs and the length differences in the SSC, the first two PCoA coordinates explained >60% of the total variance in both individuals, although the absolute variance values along the first two coordinates suggested a higher total variance for Cb01A than for Cb04B.

**Figure 3.**
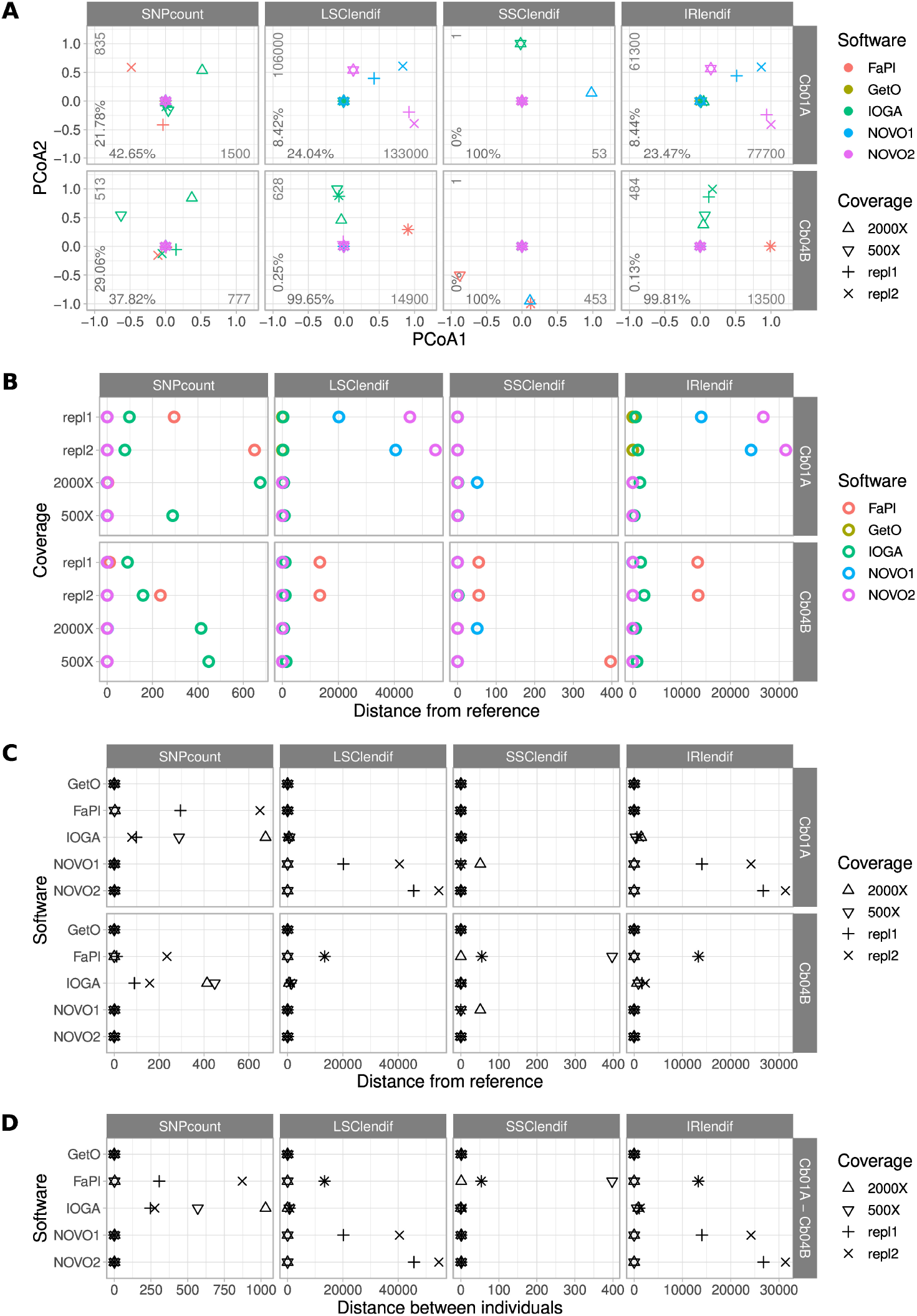
Comparisons of the number of SNPs and the lengths of each the four genome regions across the plastid genome assemblies of *C. bakuense* as generated by different assembly software tools and depths of sequence coverage. Subplot **(A)** displays the results of PCoAs, subplots **(B)** and **(C)** the results of comparisons between a target assembly and the final plastid genome sequence, and subplot **(D)** the results of assembly comparisons between the two individuals of *C. bakuense* under study. In the PCoA plots, the first and second principal coordinate are displayed, the percentages indicate the variance explained along the respective axis, and the integers express the range of the data. The abbreviations for the four distance metrics are: ‘SNPcount’ for the total number of SNPs between two assemblies; ‘LSClendif’, ‘SSClendif’, and ‘IRlendif’ for the differences in sequence length in the LSC, SSC, and IR region between two assemblies, respectively.

The comparison of pairwise genetic distances between the assemblies of different software tools highlighted the presence of SNPs between the final genome sequences and the assemblies generated with FastPlast and IOGA (Figure 3B). It also highlighted the presence of IR and LSC length differences between the final genome sequences and the assemblies generated wit FastPlast and NOVOPlasty. The overall similarity of the length difference patterns for the LSC and the IR suggests that length deviations in either region are compensated by a corresponding change in the other region during genome assembly, rather than changes of the SSC.

The comparison of pairwise genetic distances between the assemblies of different depths of sequence coverage highlighted that the observed length and sequence deviations to the final genome sequences were not constant across different sequence coverage depths, except for the assemblies generated with GetOrganelle (Figure 3C). For example, for assemblies generated with IOGA, the reduction of sequence coverage depth had an almost random effect on SNP count and region length, as it decreased the number of SNPs and the IR/LSC length difference in Cb01A when compared to the final genome sequences, whereas it increased both factors in Cb04B. GetOrganelle was the only software to produce assemblies with the same sequence and region lengths across all assembly parameters.

The comparison of genetic distances between the assemblies of the two individuals of *C. bakuense* under study demonstrated that only GetOrganelle consistently and repeatably generated the same plastid genome sequences across different software tools and depths of sequence coverage (Figure 3D). A length difference of only one bp (located in the SSC) was found to separate all assemblies produced by GetOrganelle for the two individuals. Under FastPlast and IOGA, by contrast, the number of SNPs detected between the two assemblies was far greater and sometimes even larger than the SNP differences to the respective final genome sequence. Moreover, under these software tools the number of SNPs between different assemblies of the same individual did not sum up to the number of SNPs between individuals, suggesting that at least some of the SNPs were shared between the assemblies of the same individual. Remarkably, the differences in SSC length for assemblies generated with NOVOPlasty under seed sequence 1 and a coverage cap of 2,000x were identical between the two individuals of *C. bakuense,* suggesting that the sequence difference causing this length deviation is identical in the assemblies of both individuals and probably caused by the specific seed sequence employed.

The visual comparison of the lengths of the four plastid genome regions across the different genome assemblies indicated that the differences in total genome length were primarily correlated with length changes in the LSC and the IRs (Figure 4). While the length of the SSC was virtually constant across all software tools and sequence coverage depths (~13,400 bp; Table S1 in Supplementary Material), the length of the IR was highly sensitive to the precise assembly conditions. Especially in assemblies generated with NOVOPlasty for Cb01A as well as with FastPlast for Cb04B, the IR lengths varied by a factor of 1.5 to 2, which was partially compensated by a corresponding reduction of the LSC length, sometimes to less than half of the length displayed in other assemblies. A complete list of the lengths of the four plastid genome regions in relation to the different software tools, levels of sequence coverage depth, seed sequences, and run replicates is given in Table S1 for Cb01A and Table S2 for Cb04B (Supplementary Material).

**Figure 4.**
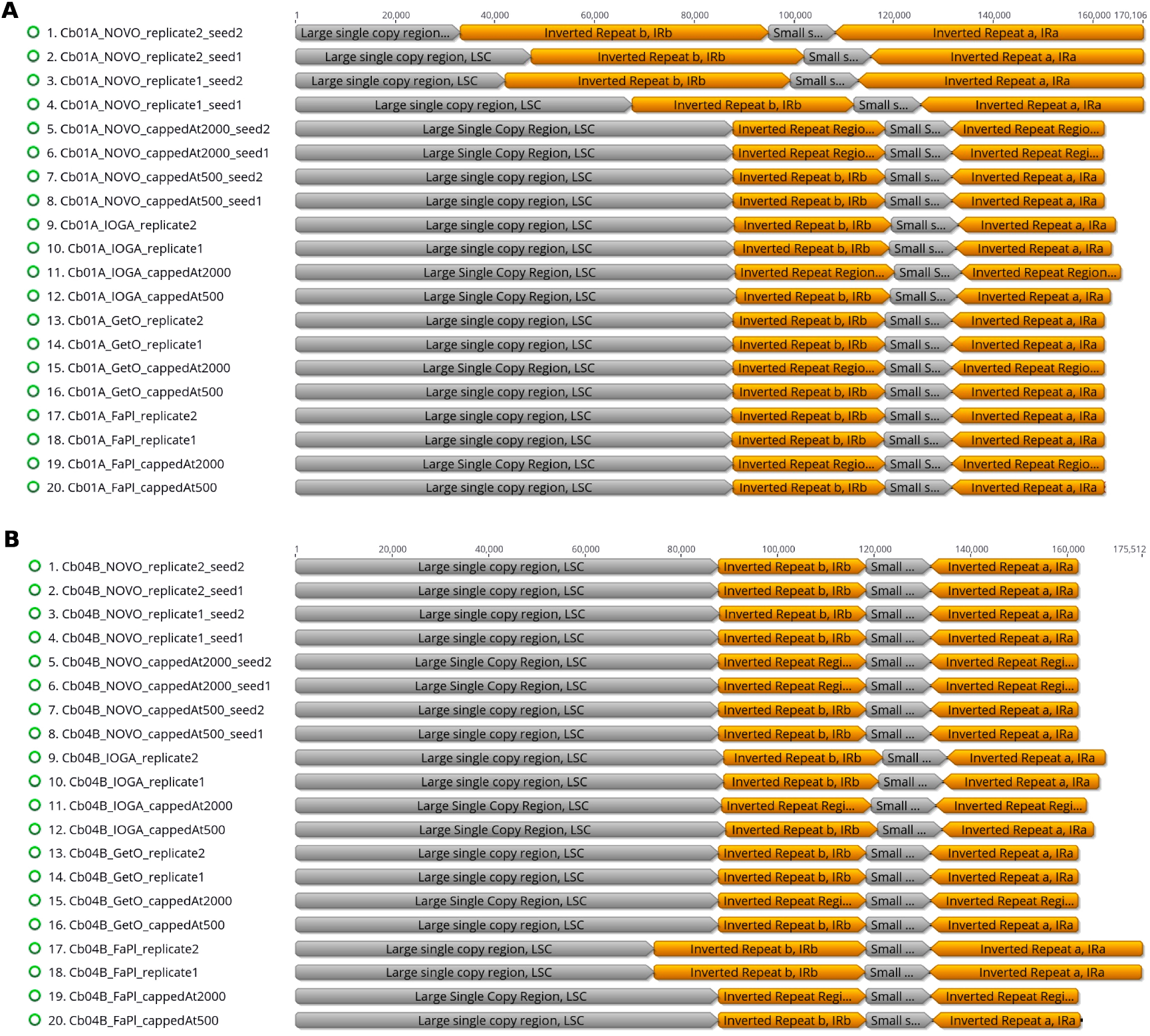
Overview of the relative lengths of the LSC, the SSC, and the two IRs across the plastid genome assemblies as generated by different assembly software tools and depths of sequence coverage. Subplot **(A)** displays the assemblies for plant individual Cb01A, subplot **(B)** for Cb04B.

### Differences in gene content and annotations

The nucleotide and length differences between the assembled plastid genomes were located in both the coding and the non-coding sections of the genomes and often manifested themselves as differences in gene content (Table 2). Specifically, the annotated sequences of several assemblies either lacked certain protein- and tRNA-coding genes due to missing genome sections or exhibited non-functional protein-coding genes due to internal stop codons caused by nucleotide polymorphisms. All assemblies generated with IOGA, for example, exhibited housekeeping genes with internal stop codons. Among the assemblies generated with FastPlast, replicate runs 1 and 2 for Cb01A and replicate run 2 for Cb04B under the original coverage depth as well as the assembly of Cb04B under a coverage cap of 500x produced gene sequences with internal stop codons. Similarly, the length differences between the four plastid genome regions across the assemblies generated with NOVOPlasty for Cb01A correlated with a lack of up to 17 different genes compared to the final genome sequence of the plant individual, even when all assembly contigs were concatenated to a super-contig; this result was observed for both seed sequences and, thus, appears to be independent of the internal start point of the genome assembly. All of the missing genome regions in the assemblies generated with NOVOPlasty were notably located at the 5’ end of the LSC, suggesting a potential bias in the assembly of this genome region. All plastid genome assemblies generated with GetOrganelle, by contrast, exhibited a complete gene complement and a full genome size.

**Table 2.** Overview of incorrect or missing annotations among the plastid genome assemblies of *C. bakuense* as generated under different assembly software tools, sequence coverage depths, seed sequences, and run replicates. All plastid genome assemblies generated with GetOrganelle for both plant individuals and with NOVOPlasty for Cb04B exhibited a complete gene complement and a full genome size and are, thus, not listed. The last column denotes cases where the DNA sequence is missing at a given position despite the assembly being circular and indicated as complete. Superscript ^a^ indicates a location in IRa, superscript ^b^ in IRb. Abbreviations used as in Table 1.

| Asmb. | Cov. | Repl. | NOVO seed | Internal stop codons in translation | No DNA sequence at this position |
| --- | --- | --- | --- | --- | --- |
| *Cb01A* |  |  |  |  |  |
| FaPl | orig. | repl1 |  | rpl23<sup>a,b</sup>, rrn16<sup>a,b</sup> |  |
| FaPl | orig. | repl2 |  | rpl2<sup>a,b</sup>, ycf2<sup>a,b</sup>, rpl23<sup>a</sup> |  |
| FaPl | 2000x |  |  |  |  |
| FaPl | 500x |  |  |  |  |
| IOGA | orig. | repl1 |  | psbA, rpl23<sup>a,b</sup>, ycf2<sup>a,b</sup>, rrn16<sup>a,b</sup> |  |
| IOGA | orig. | repl2 |  | psbA, ycf2<sup>a,b</sup> |  |
| IOGA | 2000x |  |  | psbA, ycf2<sup>a,b</sup>, ycf1<sup>a,b</sup>, ndhH |  |
| IOGA | 500x |  |  | psbA, rps2<sup>a,b</sup>, ycf2<sup>a,b</sup>, ndhH | trnH-GUG |
| NOVO | orig. | repl1 | seed1 |  | trnH-GUG, psbA, trnK-UUU, matK, rps16 |
| NOVO | orig. | repl2 | seed1 |  | trnH-GUG, psbA, trnK-UUU, matK, rps16, trnQ-UUG, psbK, psbI, trnS-GCU, trnG-UCC, trnR-UCU, atpA, atpF, atpH, atpI, rps2, rpoC2 |
| NOVO | 2000x |  | seed1 |  |  |
| NOVO | 500x |  | seed1 |  |  |
| NOVO | orig. | repl1 | seed2 |  | trnH-GUG, psbA, trnK-UUU, matK, rps16, trnQ-UUG, psbK, psbI, trnS-GCU, trnG-UCC, trnR-UCU, atpA, atpF, atpH, atpI, rps2, rpoC2 |
| NOVO | orig. | repl2 | seed2 |  | trnH-GUG, psbA, trnK-UUU, matK, rps16, trnQ-UUG, psbK, psbI, trnS-GCU, trnG-UCC, trnR-UCU, atpA, atpF, atpH, atpI, rps2, rpoC2 |
| NOVO | 2000x |  | seed2 |  |  |
| NOVO | 500x |  | seed2 |  |  |
| *Cb04B* |  |  |  |  |  |
| FaPl | orig. | repl1 |  |  |  |
| FaPl | orig. | repl2 |  | rpl23<sup>a</sup> |  |
| FaPl | 2000x |  |  |  |  |
| FaPl | 500x |  |  | ndhF |  |
| IOGA | orig. | repl1 |  | psbA, rpl23<sup>b</sup>, rpl2<sup>a</sup> |  |
| IOGA | orig. | repl2 |  | psbA, ndhH, rpl23<sup>a</sup>, rpl2<sup>a,b</sup> |  |
| IOGA | 2000x |  |  | psbA, petB |  |
| IOGA | 500x |  |  | psbA, rps23<sup>a,b</sup> |  |

### Sequence coverage and SNP location

Visualizing the location of SNPs across the genome assemblies indicated a possible association of their location with regions of low sequence coverage. The average genome-wide sequence coverage depth for the final plastid genome of Cb01A was 8,410x, the average genome-wide coverage depth for the final plastid genome of Cb04B was 5,430x. We found that some of the plastid genome assemblies generated in this study exhibited a considerable number of SNPs when compared to the final genome sequence and that these SNPs were often associated with regions of reduced sequence coverage depth. For example, the IRs of the plastid genome assemblies of Cb01A that were generated with FastPlast contained two adjacent calculation windows with a coverage depth of only 1,200x and 2,300x, respectively; these coverage depths represented only 14% and 27% of the genome-wide average coverage depth (Figure 5). The two windows were located between the tRNA genes *trnV-GAC* and *trnI-GAU* and covered parts of the gene coding for the 16S rRNA subunit *(rrn16).* Compared to the final genome sequence of Cb01A, the assemblies of both replicate runs exhibited a high density of SNPs in the very same region (Figure 5, circles A and B); SNPs outside this particular region also existed but were clustered less densely, if at all. Similarly, a high density of SNPs was found in replicate run 2 at the replication origin of the genome, which also exhibits a considerably reduced coverage depth (Figure 5, circle B); however, the reduced coverage depth at the replication origin represents an artifact introduced by the mapping software during the extraction of plastid genome reads from the raw read set and should, thus, not be seen as a region with naturally reduced coverage depth. The assemblies generated with FastPlast under the capped read sets, by contrast, did not exhibit SNPs compared to the final genome sequence (Figure 5, circles C and D). A similar interdependence between the location of SNPs and regions with reduced coverage depth was observed for the assemblies of Cb01A generated with IOGA (Figure S1 in Supplementary Material); the amount and the distribution of SNPs in comparison to the final genome sequence were, however, greater than in the assemblies with FastPlast and neither restricted to the IRs nor any particular read set. By comparison, the plastid genome assemblies generated with GetOrganelle or NOVOPlasty did not display any SNPs in comparison to the final genome sequence, irrespective of a cap on coverage depth.

**Figure 5.**
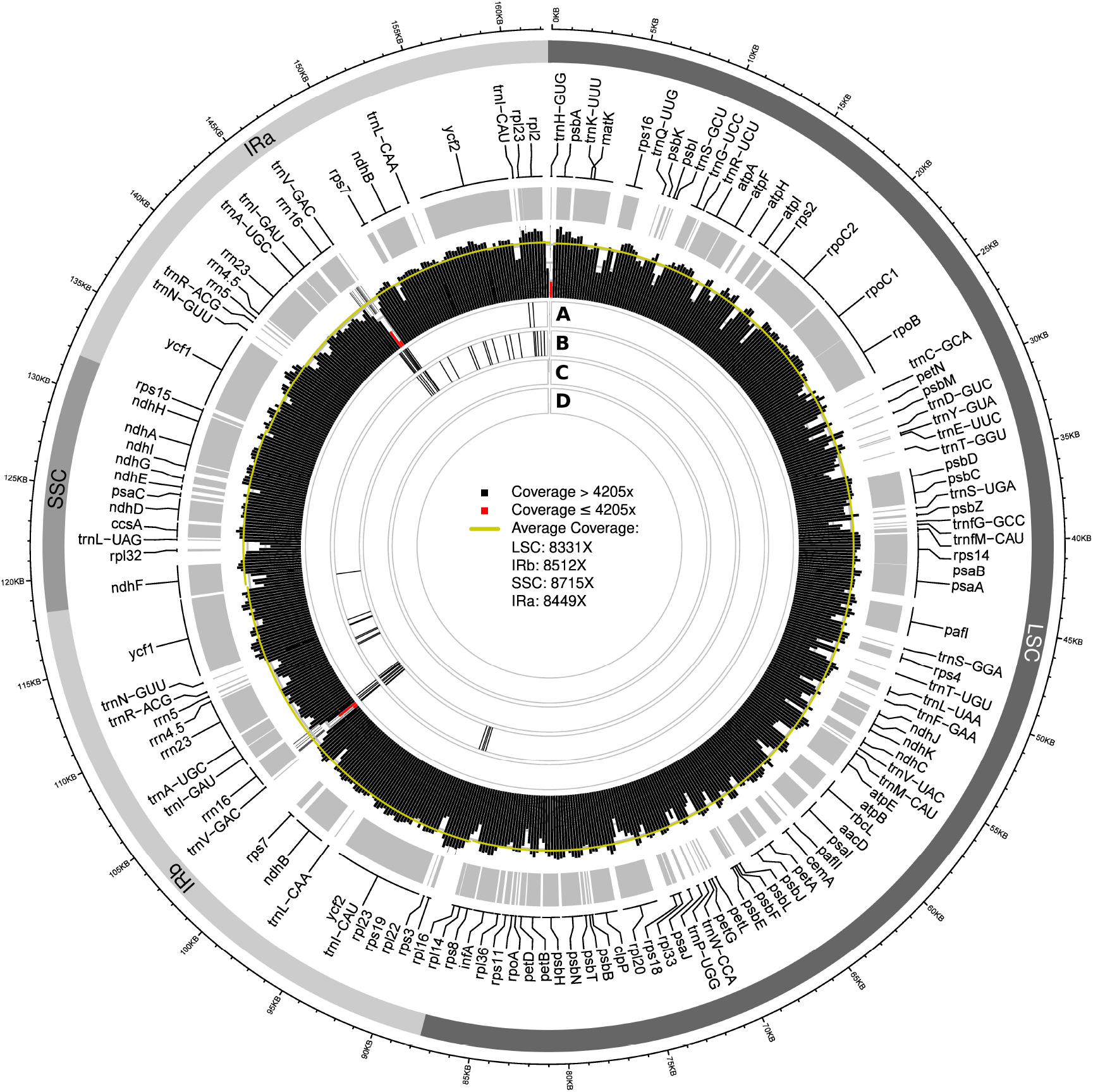
Visualization of the sequence coverage depth of the plastid genome of Cb01A generated with FastPlast relative to the location of SNPs for assemblies generated under different sequence coverage depths, the location of genes, and the quadripartite genome structure. Red bars in the visualization of sequence coverage depth indicate calculation windows with a coverage depth that is equal to, or less than, 50% of genome-wide average coverage depth. The four rings beneath the coverage depth visualization indicate the location of SNPs relative to the final genome sequence when the assembly is conducted under different run replicates and coverage caps: rings **(A)** and **(B)** display the location of SNPs in replicate runs 1 and 2, respectively, under the original coverage depth; ring **(C)** displays the location of SNPs in the assembly generated under a coverage cap of 2,000x; ring **(D)** displays the location of SNPs in the assembly generated under a coverage cap of 500x. Black bars within each ring represent the occurrence of three SNPs per 100 bp.

### Phylogenetic inference

The results of our phylogenetic tree reconstruction on the combined set of all plastid genome assemblies of *C. bakuense* generated in this study and the 21 plastid genome records of other species of *Calligonum* did not indicate that the sequence variability across the assemblies of *C. bakuense* was large enough to affect its phylogenetic placement within *Calligonum* (Figure S2 in Supplementary Material). While the different genome assemblies of *C. bakuense* did not cluster by individual, software tool, or level of sequence depth, several assemblies with a comparatively high level of SNPs in comparison to the final genome sequence (i.e., those generated with IOGA and FastPlast) formed a separate, albeit poorly supported clade, which possibly represents the result of long-branch attraction. The nucleotide differences between most of the other assemblies were not phylogenetically informative and, thus, did not result in the identification of specific clades among the assembly sequences. However, the results of this tree reconstruction indicated that the observed sequence differences among the assemblies generated with different software tools, levels of sequence coverage depth, seed sequences, and run replicates may potentially influence intra-specific evolutionary analyses of *C. bakuense*. The results of our tree reconstruction to infer the phylogenetic position of *C. bakuense* among other species of *Calligonum* recovered the final plastid genomes of *C. bakuense* as sister to *C. arborescens* (Figure 6). The sister relationship between *C. bakuense* and *C. arborescens* was weakly supported (BS 29%) but both taxa were recovered as part of a fully-supported clade alongside *C. caput-medusae*. Overall, the reconstruction recovered the same phylogenetic relationships as reported by Song et al. (2020), indicating that the inclusion of *C. bakuense* did not alter alter the tree reconstruction of the genus.

**Figure 6.**
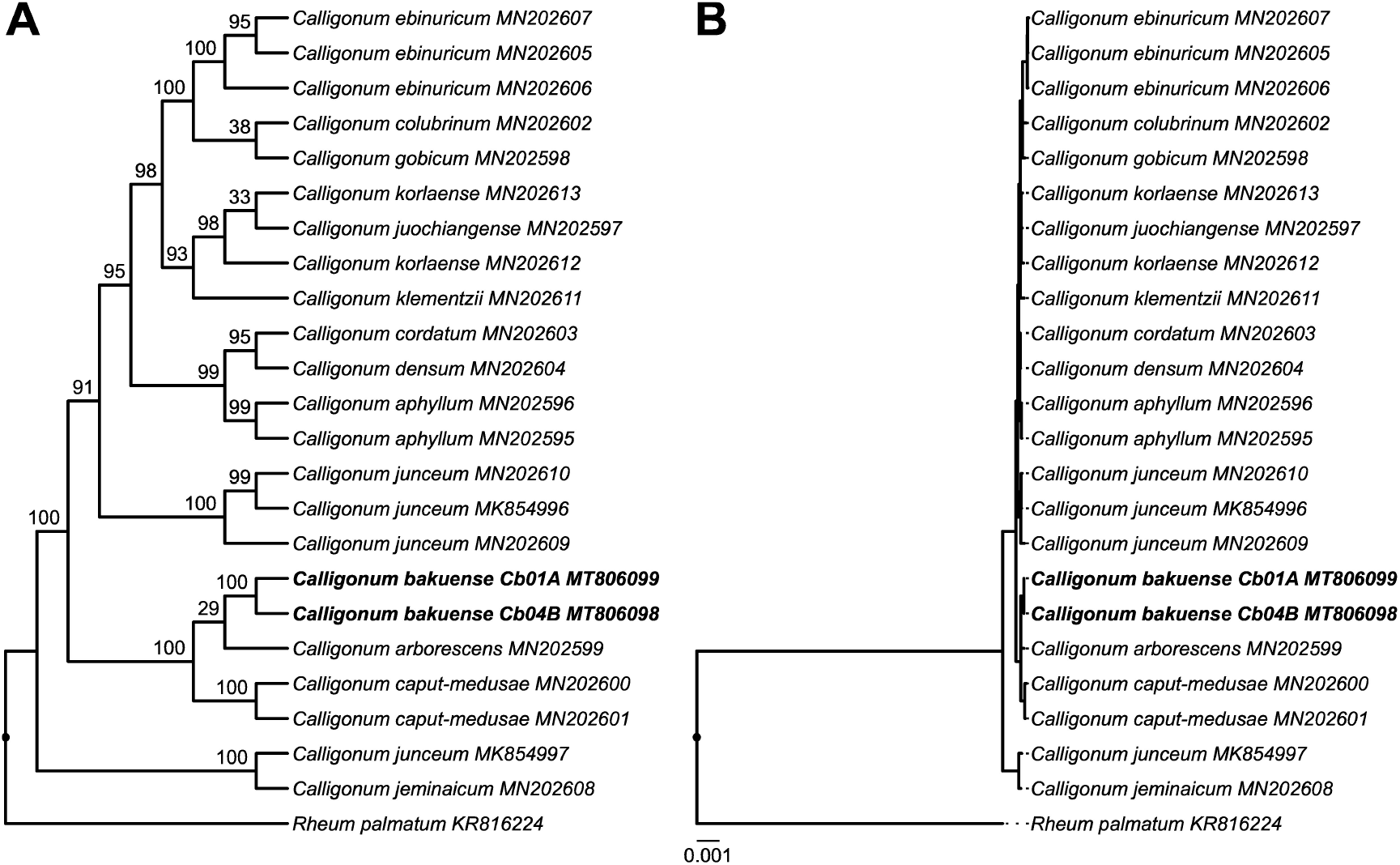
Phylogenetic position of *C. bakuense* among other species of *Calligonum. C. bakuense* is represented by the final plastid genomes of plant individuals Cb01A and Cb04B, which are highlighted in bold. The displayed phylogenetic tree represents the best tree inferred under ML, visualized as **(A)** cladogram with statistical node support and **(B)** the corresponding phylogram with exact branch lengths. BS support values are given above the branches of the cladogram.

## DISCUSSION

### Phylogenetic position of *C. bakuense* based on complete plastid genomes

This investigation is the first to report the complete plastid genome of *C. bakuense* and, thus, further advances the study of *Calligonum*, as knowledge of the plastid genome of this Caucasian endemic supports research on the evolutionary diversification of the genus. For example, our analyses underscore the potential of complete plastid genome sequences for resolving species level relationships in *Calligonum* (e.g., Song et al., 2020), whereas individual regions of the plastid genome appear to yield insufficient phylogenetic information (e.g., Tavakkoli et al., 2010). The results of our phylogenetic analyses (Figure 6) only partially agree with the current taxonomic classification of *Calligonum*. *Calligonum bakuense* is a member of sect. *Calligonum*, yet was recovered as part of a clade formed by individuals of *C. arborescens* and *C. caput-medusae,* both of which are members of sect. *Medusa* Sosk. et Alexandr (Soskov, 2011). The current sectional classification of *Calligonum* is primarily based on differences in fruit morphology and, as suggested by Song et al. (2020), probably not natural, and our results provide further evidence for this. The phylogenetic position of *C. bakuense* alongside *C. arborescens* and *C. caput-medusae* is, however, supported on the basis of their geographic distribution. *Calligonum arborescens* and *C. caput-medusae* are both distributed in steppes eastwards of the Caspian Sea, ranging from Turkmenistan to China. Based on similarities in fruit morphology and the tetraploid nature of *C. bakuense* (2n=36; Bolkhovskikh et al., 1969), the species was hypothesized to be an allotetraploid that arose from ancestors of *C. polygonoides* L. and *C. acanthopterum* I.G. BORSHCH (Soskov and Akhmed-Zade, 1974). While *C. polygonoides* is widespread and also occurs in Azerbaijan (Karjagin, 1952), *C. acanthopterum* is known only from Kazakhstan and Turkmenistan. By contrast, the widespread species *C. aphyllum*, which is distributed from North Africa to the Caucasus (including Azerbaijan) and China, is morphologically distinct from *C. bakuense* (e.g., winged fruits that lack bristles) and probably not a close relative to *C. bakuense*; the results of our phylogenetic analyses support this interpretation. Future phylogenetic investigations should, thus, increase both the taxon sampling and, where possible, the geographic representation of the more widespread taxa of *Calligonum* such as *C. polygonoides*. The inclusion of the non-coding sections of the plastid genome (i.e., introns and intergenic spacers) in a genus-wide plastid phylogenomic analysis of *Calligonum* represents another task for future phylogenetic studies on the genus; such analyses were found to be essential in clarifying the phylogenetic history of other angiosperm genera with low genetic distances among species (e.g., *Gynoxys;* Escobari et al., 2021). The inclusion of phylogenetic information from the nuclear genome will also be relevant for clarifying the phylogenetic history of *Calligonum,* especially regarding possible reticulate speciation events within the genus.

The single basepair difference detected between the plastid genomes of two individuals of *C. bakuense* falls within the spectrum of sequence divergence for plastid genomes of narrow endemic plant species. While intra-specific comparisons of complete plastid genomes are rare (e.g., Jiang et al., 2017; Teshome et al., 2020), the investigations that do exist often report of only a handful of SNPs between plant individuals. The narrow endemic *Pinus torreyana*, for example, had five SNPs between the plastid genomes of two individuals from both parts of its disjunct distribution range (Whittall et al., 2010, indels not reported). Similarly, at least two SNPs and one indel were found between two plastid genomes of *Fagus multinervis* endemic to Ulleung Island off the South Korean coast (Yang et al., 2020). However, it is not possible to evaluate the genetic diversity of the species or any well-founded conservation genetic assessments based on the sequences of two plastid genomes alone. Instead, more research on the genetic diversity within *C. bakuense*, including the use of nuclear microsatellites, is needed.

### Impact of software choice on plastid genome assembly

By comparing the assembly contigs of *C. bakuense* that were generated with four different assembly software tools, we found that software choice can have an inordinate influence on the sequence and structure of plastid genome sequences and that the assembly results of some these tools need to be treated with caution. Among the differences across the assemblies were the presence of SNPs (compared to the final genome sequences), the loss of entire genes or their functionality, and the expansion and contraction of the IRs (as well as compensatory length changes in the LSC). Such occurrences have been occasionally interpreted in an evolutionary context (e.g., Mohanta et al., 2020), but it stands to reason that at least some of the differences between the plastid genomes of closely related species may have a more technical origin, as recently demonstrated by Freudenthal et al. (2020). The results of our investigation support the hypothesis that differences among plastid genome assemblies may also be technical in nature: we found that the plastid genome assemblies of different software tools exhibited considerably different genome sequences despite employing the same input data and that some of the assembly contigs could not be replicated in different runs of the same software (Table 1). Only the software GetOrganelle was found to generate consistent and repeatable results for both datasets. FastPlast, by contrast, was found to be prone to the introduction of SNPs and, in some cases, also structural deviation among the assemblies. NOVOPlasty introduced few, if any, SNPs compared to the final genome sequences but exhibited a tendency for generating structural deviations, which even occurred when the same assembly was conducted with different seed sequences. The assemblies generated with IOGA were fragmentary in all cases and exhibited numerous SNPs and structural deviations compared to the final genome sequences. Worse still, we found that many of these sequence deviations generated by IOGA would result in incorrect conclusions about gene content and functionality when compared to the final genome sequences (Table 2). We, therefore, concur with Freudenthal et al. (2020) that users should abstain from employing the software IOGA (which is no longer maintained) for plastid genome assemblies, and that the software tools FastPlast and NOVOPlasty should be employed with caution and only in cases where GetOrganelle fails to produce assembly contigs. We also concur with the suggestion that the replication of assembly results across different software runs and seed sequences (where applicable) are beneficial precautions in the generation of trustworthy plastid genome sequences.

Our results do not imply that the assemblies generated with GetOrganelle necessarily represent the true plastid genome sequences. It is theoretically possible for a software tool to consistently and repeatably produce incorrect results, and we also cannot rule out the presence of more than one unique plastid genome per plant individual (Scarcelli et al., 2016; Wang and Lanfear, 2019). However, the software tools FastPlast and NOVOPlasty produced the same genome sequence as identified through GetOrganelle under some of the evaluated settings. We, therefore, considered the plastid genome assemblies generated with GetOrganelle under the read sets capped at a coverage depth of 500x as the most likely genome sequences for the two individuals of *C. bakuense* and employed them as the final plastid genomes. Aside from the idiosyncrasies introduced by different assembly software, the observed differences among the plastid genome assemblies may also be the result of nucleotide polymorphism among the input reads (Scarcelli et al., 2016). Such polymorphism within the read set could represent genuinely different variants of the plastid genome (i.e., heteroplasmy; Walker et al., 2015; Wang and Lanfear, 2019), genomic transfers of sections of the plastid to the nuclear or the mitochondrial genome, followed by a pseudogenization of the transferred regions (Ruhlman and Jansen, 2014), or sequencing errors during data generation (Nakamura et al., 2011), and may be decoded differently by the different assembly software.

### Impact of sequence coverage depth on plastid genome assembly

By comparing the assembly contigs of *C. bakuense* that were generated under three different levels of sequence coverage depth, we found that sequence coverage can also have an important impact on plastid genome assembly. Specifically, we found that the capping of sequence coverage prior to genome assembly had a measurable effect on the number of assembly contigs constructed, the nucleotide sequences of these contigs, the length of the different plastid genome regions (particularly the IRs), and the number of valid gene annotations. The effects of capping coverage depth were measurable in Cb01A but especially pronounced in Cb04B and suggested the trend that a cap in coverage depth rendered the assemblies more consistent in sequence and length. Specifically, the reduction of coverage depth appeared to improve the replicability of the genome assemblies under FastPlast and NOVOPlasty (Table 1), but this effect may rather represent the concomitant increase in coverage evenness. Both the original and the capped read sets of the two individuals of *C. bakuense* analyzed here vastly exceed the recommended coverage depth for plastid genome assembly (Twyford and Ness, 2017; McKain et al., 2018). For example, with an average genome-wide coverage depth of 8,410x (for the final plastid genome), and a minimal coverage depth of more than 1,000x in any genome position, the original read set of Cb01A comprises more than enough sequence information to completely assemble the plastid genome. The failure to assemble the genome with some of the tested software tools is, thus, more likely associated with coverage unevenness than coverage depth. A high but comparatively even depth of sequence coverage may be the best strategy for a successful plastid genome assembly under the tested software tools.

Our results are congruent with the findings of other investigations that report an impact of sequence coverage on the genome assembly process (Pedersen et al., 2017) or a correlation between local extremes in sequence coverage and assembly contig deviations (Kim et al., 2015). In general, the depth of sequence coverage is indicative for a reliable identification of sequence rearrangements and other structural variants (Izan et al., 2017; Sims et al., 2014), but the relationship between coverage depth and assembly reliability is not straightforward. While greater sequence coverage typically increases the chance that rearrangement endpoints are captured and confirmed by multiple reads (Chen et al., 2009), genomic regions with exceptionally high depth of sequence coverage have also been reported as problematic for the identification of SNPs (Li, 2014).

### Impact of assembly differences on phylogenetic placement

The results of this investigation illustrate that the correct assembly of plastid genomes without a subsequent evaluation cannot be taken for granted, even with dedicated software tools. Incorrect genome assemblies have the potential to affect downstream analyses, such as studies on genetic diversity or evolutionary history. Even if the assembly differences observed here did not seem to affect the inferred phylogenetic position of *C. bakuense* within *Calligonum* under the current taxon sampling, we cannot exclude that errors introduced by the assembly software can lead to incorrect phylogenetic reconstructions, especially if samples with low genetic distances are analyzed.

### Recommendations for future studies

Given the results of this investigation, we conclude three recommendations for the application of *de novo* plastid genome assembly. First, we recommend to compare the assembly results of different software tools and multiple software runs before accepting any assembly as the final genome sequence. As demonstrated here, different assembly software may vary considerably in their accuracy and repeatability. We, therefore, recommend to consider only such results for subsequent analysis that are reproducible across different software tools and replicate runs. Users are hereby not restricted to the four software tools tested in this investigation: there are various software applications for the *de novo* assembly of complete plastid genomes from genome skimming data, including general short-read assemblers. Among such general assemblers are SOAPdenovo2 (Luo et al., 2012), Platanus (Kajitani et al., 2014), and Meraculous (Chapman et al., 2011), which we tested on *C. bakuense* in a preliminary analysis of this investigation. However, we found that for *C. bakuense* only Platanus generated assembly contigs that represented either the complete plastid genome (Cb01A) or sections of it (Cb04B). Second, we recommend to cap the sequence coverage of the input read data to an approximately even distribution along the whole genome sequence while keeping the overall coverage depth high before conducting plastid genome assembly. While the exact relationship between assembly accuracy and both coverage depth and coverage evenness is poorly understood for plastid genome assembly, the results of such investigations on bacterial genomes indicate a considerable impact of both factors (Magoc et al., 2013; Pedersen et al., 2017). More research is needed to determine the optimal balance between coverage depth and coverage evenness for reliable plastid genome assembly. Third, we recommend the release of the raw sequence data and detailed information on the genome assembly and annotation process during the publication of new plastid genomes. Only by sharing the input data as well as a precise description of the type and sequence of the software tools employed are assembly results genuinely reproducible and, ultimately, reliable (Gruening et al., 2018; Gruenstaeudl et al., 2018). The provisioning of the raw sequence data as well as detailed assembly and annotation information is also essential if researchers wish to re-analyze the data with new and improved methods (e.g., Gruenstaeudl, 2019). Expressly for this purpose we release the raw sequence reads, the read datasets capped at different coverage depths, and the raw assembly results from this study to the public.

## DECLARATIONS

### Author Contributions

The study was devised by KR and MG, with participation from TB and VK. The distribution data were assessed by VK and visualized by KR. Preliminary data analyses were conducted by CC and MG, final analyses by EG and MG. EG performed all post-assembly finishing steps, MG performed the sequence comparisons and the phylogenetic reconstructions, and KR calculated the PCoAs. The writing of the manuscript was lead by EG, MG, and KR, with additional input by TB. All authors have read and approved the final version of the manuscript.

## Acknowledgments

We thank Gerald Parolly, Nadja Korotkova, and Tural Qasimov for assistance with field collection, Bettina Giesicke for DNA extraction, Halil Atis for Illumina library preparation, and Cathrin Schierenbeck for assistance with sequence data archiving. The authors acknowledge the Berlin Center of Genomics in Biodiversity Research for providing lab assistance and the high-performance computing service of the ZEDAT of the Freie Universität Berlin for providing allocations of computing time. We also acknowledge support by the Open Access Publication Initiative of Freie Universitat Berlin. Several of the analyses presented here represent part of a thesis by EG toward a master of science degree.

## Data Availability Statement

The datasets generated for this study can be found on the NCBI Sequence Read Archive (https://www.ncbi.nlm.nih.gov/sra) under identifiers SRX9433946 (Cb01A) and SRX9433941 (Cb04B). The datasets analyzed for this study can be found on Zenodo under record number 3999863 (https://zenodo.org/record/3999863).

## Supplemental Data

**Table S1** | Numeric comparison of the plastid genome assemblies of plant individual Cb01A as generated by different software tools, levels of sequence coverage depth, seed sequences, and run replicates.

**Table S2** | Numeric comparison of the plastid genome assemblies of plant individual Cb04B as generated by different software tools, levels of sequence coverage depth, seed sequences, and run replicates.

**Figure S1** | Visualization of the sequence coverage depth of the plastid genome of Cb01A generated with IOGA relative to the location of SNPs for assemblies generated under different sequence coverage depths, the location of genes, and the quadripartite genome structure.

**Figure S2** | The phylogenetic position of all plastid genome assemblies of *C. bakuense* as generated with different software tools, levels of sequence coverage depth, seed sequences, and run replicates in relation to other species of *Calligonum.*

