## Supplemental Materials for "Software choice and depth of sequence coverage can impact plastid genome assembly – A case study in the narrow endemic *Calligonum bakuense*"

**Table S1:** Numeric comparison of the plastid genome assemblies of plant individual Cb01A as generated by different software tools, levels of sequence coverage depth, seed sequences, and run replicates. Numbers in parentheses indicate numbers as duplicated in the IR.

| | FaPl 2000X | FaPl 500X | FaPl repl1 | FaPl repl2 | GetO 2000X | GetO 500X | GetO repl1 | GetO repl2 | IOGA 2000X | IOGA 500X |
|---|---|---|---|---|---|---|---|---|---|---|
| Genome size (bp) | 162,128 | 163,006 | 162,093 | 162,173 | 162,128 | 162,128 | 162,128 | 162,128 | 165,560 | 163,478 |
| LSC length (bp) | 87,689 | 87,689 | 87,642 | 87,642 | 87,689 | 87,689 | 87,689 | 87,689 | 88,194 | 88,394 |
| SSC length (bp) | 13,387 | 13,387 | 13,387 | 13,387 | 13,387 | 13,387 | 13,387 | 13,387 | 13,388 | 13,388 |
| IR length (bp) | 30,526 | 30,526 | 30,532 | 30,572 | 30,526 | 30,526 | 30,526 | 30,526 | 31,989 | 30,848 |
| Number of genes | 113 | 113 | 111 | 113 | 113 | 113 | 113 | 113 | 113 | 112 |
| Number of protein-coding genes | 79 (7) | 79 (9) | 78 (6) | 79 (7) | 79 (7) | 79 (7) | 79 (7) | 79 (7) | 79 (7) | 79 (7) |
| Number of tRNA genes | 30 (7) | 30 (7) | 30 (7) | 30 (7) | 30 (7) | 30 (7) | 30 (7) | 30 (7) | 30 (6) | 29 (7) |
| Number of rRNA genes | 4 (4) | 4 (4) | 3 (3) | 4 (4) | 4 (4) | 4 (4) | 4 (4) | 4 (4) | 4 (4) | 4 (4) |
| Number of genes with one intron (two introns) | 15 (3) | 15 (3) | 15 (3) | 15 (3) | 15 (3) | 15 (3) | 15 (3) | 15 (3) | 15 (3) | 15 (3) |
| Proportion of coding to non-coding regions | 0.71 | 0.71 | 0.68 | 0.71 | 0.71 | 0.71 | 0.70 | 0.71 | 0.69 | 0.71 |
| Average gene density (genes/kb) | 0.81 | 0.82 | 0.78 | 0.81 | 0.81 | 0.81 | 0.81 | 0.81 | 0.79 | 0.80 |
| GC content (%) | 37.5 | 37.5 | 37.4 | 37.3 | 37.5 | 37.5 | 37.5 | 37.5 | 37.3 | 37.3 |

| | IOGA repl1 | IOGA repl2 | NOVO 2000X seed1 | NOVO 2000X seed2 | NOVO 500X seed1 | NOVO 500X seed2 | NOVO repl1 seed1 | NOVO repl1 seed2 | NOVO repl2 seed1 | NOVO repl2 seed2 |
|---|---|---|---|---|---|---|---|---|---|---|
| Genome size (bp) | 163,538 | 164,532 | 162,263 | 162,128 | 162,151 | 162,128 | 170,009 | 170,003 | 170,099 | 170,106 |
| LSC length (bp) | 87,957 | 87,957 | 87,689 | 87,689 | 87,689 | 87,689 | 67,504 | 42,100 | 47,218 | 32,997 |
| SSC length (bp) | 13,387 | 13,387 | 13,336 | 13,387 | 13,387 | 13,387 | 13,387 | 13,387 | 13,387 | 13,387 |
| IR length (bp) | 31,097 | 31,594 | 30,526 | 30,526 | 30,526 | 30,526 | 44,559 | 57,258 | 54,747 | 61,861 |
| Number of genes | 109 | 114 | 113 | 113 | 113 | 113 | 108 | 96 | 99 | 95 |
| Number of protein-coding genes | 76 (5) | 79 (9) | 79 (7) | 79 (7) | 79 (7) | 79 (7) | 76 (23) | 68 (38) | 71 (34) | 67 (38) |
| Number of tRNA genes | 30 (7) | 30 (7) | 30 (7) | 30 (7) | 30 (7) | 30 (7) | 28 (7) | 24 (9) | 24 (9) | 24 (9) |
| Number of rRNA genes | 3 (3) | 5 (3) | 4 (4) | 4 (4) | 4 (4) | 4 (4) | 4 (4) | 4 (4) | 4 (4) | 4 (4) |
| Number of genes with one intron (two introns) | 15 (3) | 15 (3) | 15 (3) | 15 (3) | 15 (3) | 15 (3) | 13 (3) | 11 (3) | 12 (3) | 10 (3) |
| Proportion of coding to non-coding regions | 0.59 | 0.71 | 0.71 | 0.71 | 0.71 | 0.71 | 0.69 | 0.68 | 0.71 | 0.68 |
| Average gene density (genes/kb) | 0.76 | 0.81 | 0.81 | 0.81 | 0.81 | 0.81 | 0.84 | 0.86 | 0.86 | 0.86 |
| GC content (%) | 37.5 | 37.5 | 37.5 | 37.5 | 37.5 | 37.5 | 37.5 | 37.5 | 37.5 | 37.5 |

**Table S2:** Numeric comparison of the plastid genome assemblies of plant individual Cb04B as generated by different software tools, levels of sequence coverage depth, seed sequences, and run replicates. Numbers in parentheses indicate numbers as duplicated in the IR.

| Treatment | FaPl 2000X | FaPl 500X | FaPl repl1 | FaPl repl2 | GetO 2000X | GetO 500X | GetO repl1 | GetO repl2 | IOGA 2000X | IOGA 500X |
|---|---|---|---|---|---|---|---|---|---|---|
| Genome size (bp) | 162,129 | 163,292 | 175,272 | 175,512 | 162,129 | 162,129 | 162,129 | 162,129 | 163,822 | 165,285 |
| LSC length (bp) | 87,689 | 87,689 | 74,285 | 74,295 | 87,689 | 87,689 | 87,689 | 87,689 | 88,204 | 89,094 |
| SSC length (bp) | 13,388 | 13,785 | 13,333 | 13,333 | 13,388 | 13,388 | 13,388 | 13,388 | 13,388 | 13,388 |
| IR length (bp) | 30,526 | 30,526 | 43,827 | 43,942 | 30,526 | 30,526 | 30,526 | 30,526 | 31,115 | 31,406 |
| Number of genes | 113 | 113 | 113 | 114 | 113 | 113 | 113 | 113 | 114 | 113 |
| Number of protein-coding genes | 79 (7) | 79 (10) | 79 (23) | 80 (22) | 79 (7) | 79 (7) | 79 (7) | 79 (7) | 79 (7) | 79 (7) |
| Number of tRNA genes | 30 (7) | 30 (7) | 30 (7) | 30 (7) | 30 (7) | 30 (7) | 30 (7) | 30 (7) | 31 (6) | 30 (7) |
| Number of rRNA genes | 4 (4) | 4 (4) | 4 (4) | 4 (4) | 4 (4) | 4 (4) | 4 (4) | 4 (4) | 4 (4) | 4 (4) |
| Number of genes with one intron (two introns) | 15 (3) | 15 (3) | 15 (3) | 15 (3) | 15 (3) | 15 (3) | 15 (3) | 15 (3) | 15 (3) | 15 (3) |
| Proportion of coding to non-coding regions | 0.71 | 0.71 | 0.70 | 0.72 | 0.71 | 0.71 | 0.70 | 0.71 | 0.71 | 0.71 |
| Average gene density (genes/kb) | 0.81 | 0.82 | 0.84 | 0.84 | 0.81 | 0.81 | 0.81 | 0.81 | 0.80 | 0.79 |
| GC content (%) | 37.5 | 37.5 | 37.4 | 37.3 | 37.5 | 37.5 | 37.5 | 37.5 | 37.3 | 37.1 |

| Treatment | IOGA repl1 | IOGA repl2 | NOVO 2000X seed1 | NOVO 2000X seed2 | NOVO 500X seed1 | NOVO 500X seed2 | NOVO repl1 seed1 | NOVO repl1 seed2 | NOVO repl2 seed1 | NOVO repl2 seed2 |
|---|---|---|---|---|---|---|---|---|---|---|
| Genome size (bp) | 166,405 | 167,837 | 162,078 | 162,129 | 162,129 | 162,129 | 162,129 | 162,129 | 162,129 | 162,129 |
| LSC length (bp) | 88,717 | 88,737 | 87,689 | 87,689 | 87,689 | 87,689 | 87,689 | 87,789 | 87,689 | 87,689 |
| SSC length (bp) | 13,388 | 13,390 | 13,337 | 13,388 | 13,388 | 13,388 | 13,388 | 13,388 | 13,388 | 13,388 |
| IR length (bp) | 32,150 | 32,855 | 30,526 | 30,526 | 30,526 | 30,526 | 30,526 | 30,476 | 30,526 | 30,526 |
| Number of genes | 112 | 113 | 113 | 113 | 112 | 113 | 113 | 113 | 113 | 114 |
| Number of protein-coding genes | 78 (5) | 79 (8) | 79 (7) | 79 (7) | 78 (8) | 79 (7) | 79 (7) | 79 (7) | 79 (7) | 80 (6) |
| Number of tRNA genes | 30 (7) | 30 (6) | 30 (7) | 30 (7) | 30 (7) | 30 (7) | 30 (7) | 30 (7) | 30 (7) | 30 (7) |
| Number of rRNA genes | 4 (4) | 4 (4) | 4 (4) | 4 (4) | 4 (4) | 4 (4) | 4 (4) | 4 (4) | 4 (4) | 4 (4) |
| Number of genes with one intron (two introns) | 15 (3) | 15 (3) | 15 (3) | 15 (3) | 14 (3) | 15 (3) | 15 (3) | 15 (3) | 15 (3) | 15 (4) |
| Proportion of coding to non-coding regions | 0.66 | 0.69 | 0.71 | 0.71 | 0.71 | 0.71 | 0.70 | 0.70 | 0.71 | 0.71 |
| Average gene density (genes/kb) | 0.77 | 0.78 | 0.81 | 0.81 | 0.81 | 0.81 | 0.81 | 0.81 | 0.81 | 0.81 |
| GC content (%) | 37.5 | 37.4 | 37.5 | 37.5 | 37.5 | 37.5 | 37.5 | 37.5 | 37.5 | 37.5 |

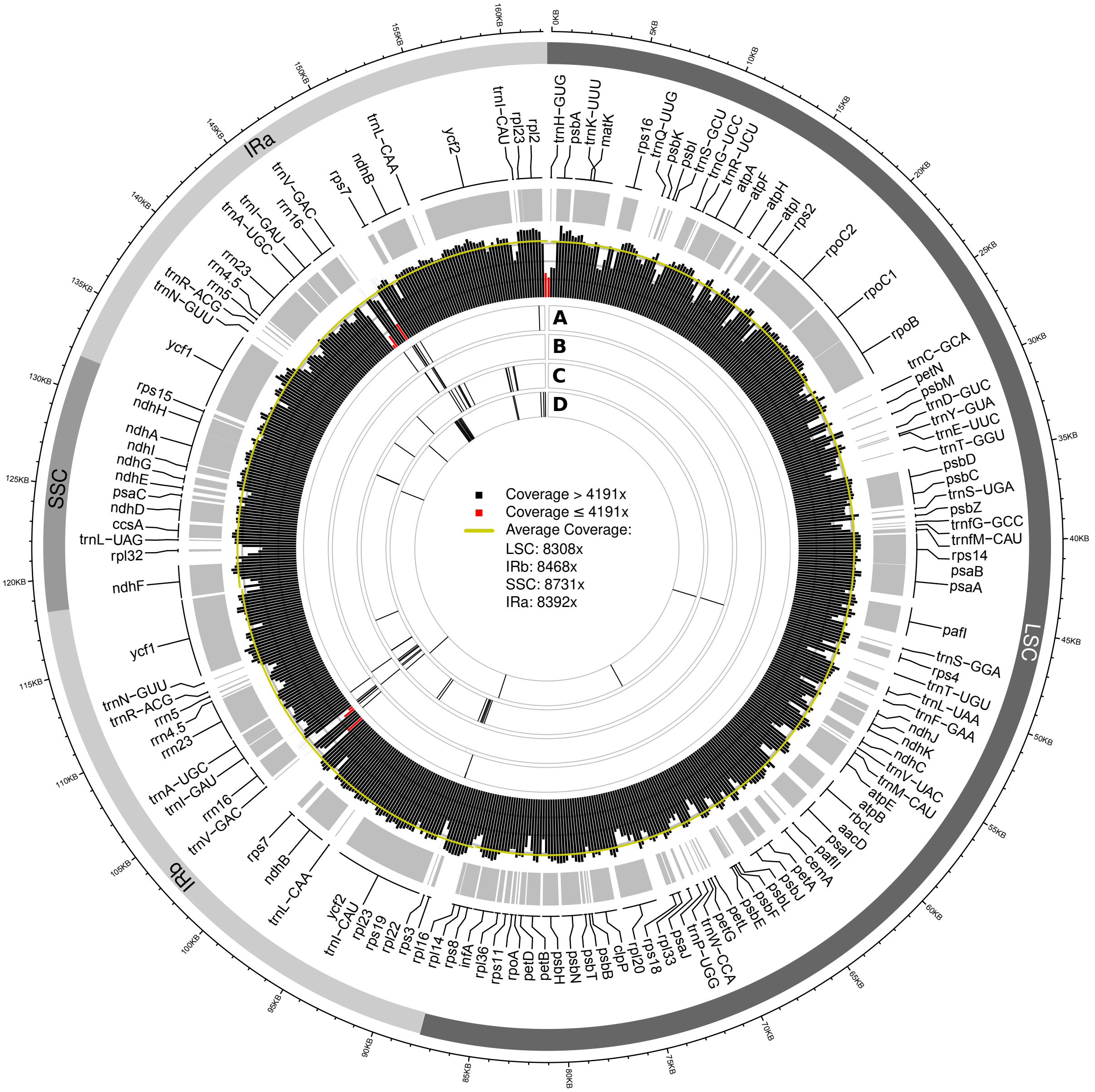


**Figure S1:** Visualization of the sequence coverage depth of the plastid genome of Cb01A generated with IOGA relative to the location of SNPs for assemblies generated under different sequence coverage depths, the location of genes, and the quadripartite genome structure. All colors, references and abbreviations are as in Figure 5 of the main text.

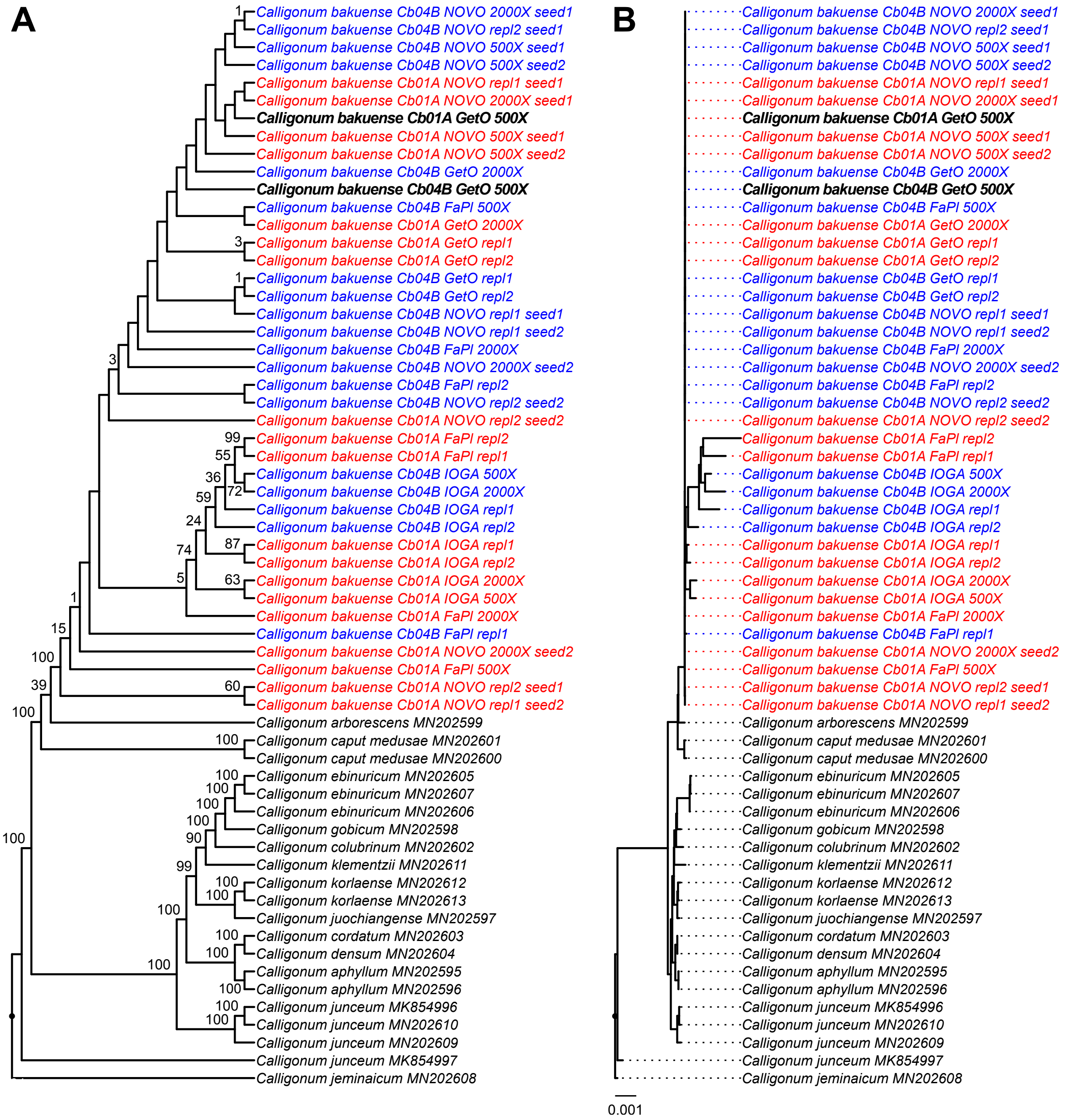


**Figure S2:** The phylogenetic position of all plastid genome assemblies of *C. bakuense* as generated with different software tools, levels of sequence coverage depth, seed sequences, and run replicates in relation to other species of *Calligonum*. Assemblies of plant individual Cb01A are highlighted in red, assemblies of Cb04B in blue. The two final plastid genome sequences of *C. bakuense* are highlighted in bold. The displayed phylogenetic tree represents the best tree inferred under ML, visualized as **(A)** cladogram with statistical node support and **(B)** the corresponding phylogram with exact branch lengths. BS support values are given above the branches of the cladogram.
